# Pandemic *Vibrio cholerae* Acquired Competitive Traits from an Environmental *Vibrio* Species

**DOI:** 10.1101/2021.05.28.446156

**Authors:** Francis J. Santoriello, Paul C. Kirchberger, Yann Boucher, Stefan Pukatzki

## Abstract

**Background:** *Vibrio cholerae,* the causative agent of cholera, is a human pathogen that thrives in estuarine environments. *V. cholerae* competes with neighboring microbes by the contact-dependent translocation of toxic effectors with the type VI secretion system (T6SS). Effector types are highly variable across *V. cholerae* strains, but all pandemic isolates encode the same set of distinct effectors. It is possible that acquisition of these effectors via horizontal gene transfer played a role in the development of pandemic *V. cholerae.*

**Results:** We assessed the distribution of *V. cholerae* T6SS loci across multiple *Vibrio* species. We showed that the fish-pathogen *V. anguillarum* encodes all three *V. cholerae* core loci as well as two of the four additional auxiliary clusters. We further demonstrated that *V. anguillarum* shares T6SS effectors with *V. cholerae,* including every pandemic-associated *V. cholerae* effector. We identified a novel T6SS cluster (Accessory Aux1) that is widespread in *V. anguillarum* and encodes the pandemic *V. cholerae* effector TseL. We highlighted potential gene transfer events of Accessory Aux1 from *V. anguillarum* to *V. cholerae.* Finally, we showed that TseL from *V. cholerae* can be neutralized by the *V. anguillarum* Accessory Aux1 immunity protein and vice versa, indicating *V. anguillarum* as the donor of *tseL* to the *V. cholerae* species.

**Conclusions:** *V. anguillarum* constitutes an environmental reservoir of pandemic-associated *V. cholerae* T6SS effectors. *V. anguillarum* and *V. cholerae* likely share an environmental niche, compete, and exchange T6SS effectors. Further, our findings highlight the fish as a potential reservoir of pandemic *V. cholerae.*

## BACKGROUND

Bacteria live in constant contact with shifting populations of bacterial competitors and predatory eukaryotic cells. Thus, effective niche colonization and survival is often dependent upon the dynamic acquisition of defense mechanisms. One such defense system is the Type VI Secretion System (T6SS), a harpoon-like nanomachine encoded by approximately 25% of all gram-negative bacteria [1–3]. The T6SS is evolutionarily related to the contractile tail of a T4 bacteriophage [4–7] and is used for the contact-dependent translocation of proteinaceous effectors into neighboring competitor cells (Fig. 1 a). Effector proteins can be toxic to non-kin bacteria and eukaryotes [1,8,9]. In the case of bactericidal effectors, the effector-secreting cell also encodes a cognate immunity protein to neutralize the effectors killing capacity and protect against attacks from sister cells [10–15]. While this secretion system is functionally conserved across gram-negative species, the core components of the T6SS vary. T6SSs can be phylogenetically classified into four types (T6SS^i^, T6SS^ii^, T6SS^iii^, and T6SS^iv^) based on multiple independent acquisition events from phages, with T6SS^i^ further divided into six subtypes (i1, i2, i3, i4a, i4b, and i5) [3,16–18].

**Figure 1.**
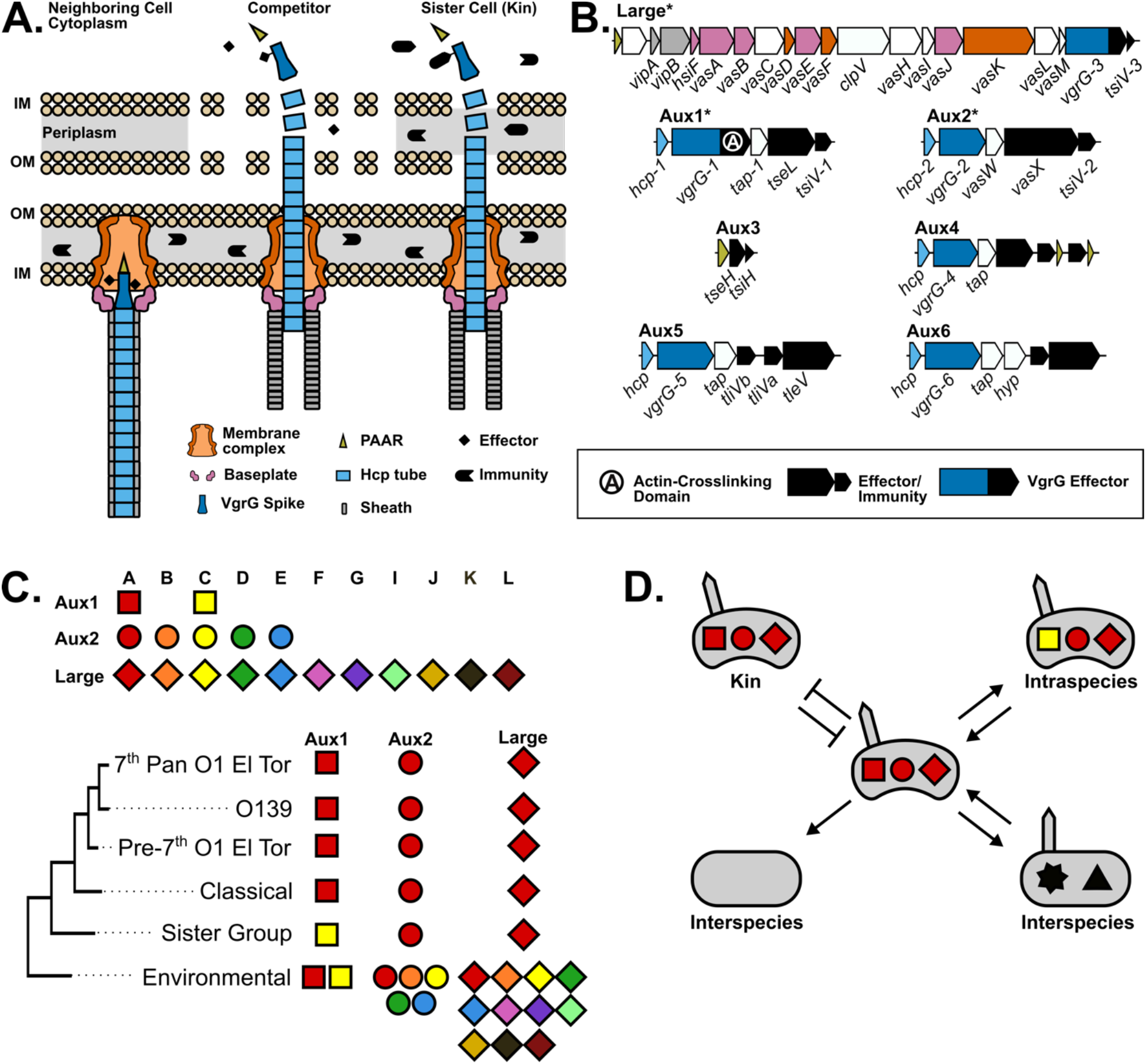
Pandemic *V. cholerae* strains encode a defined T6SS effector set for intra- and interspecies competition. **a** Schematic of the T6SS in its extended (left) and contracted (middle, right) states in competition with a neighboring non-kin (middle) or kin (right) cell. **b** Genetic diagrams of all known *Vch* T6SS loci. Coding regions are colored according to their corresponding T6SS component in (a). White coding regions are not represented in (a). Core T6SS loci are marked with an *. **c** Known *V. cholerae* effector types and their distribution across different *V. cholerae* clades. **d** Theoretical schematic of T6SS-based competition for cells with identical and differential T6SS effector sets.

The T6SS was first identified in the main pathogenic lineage of the *Vibrio cholerae* species [1], and has since been shown to be highly conserved in the *Vibrio* genus [19–24]. All *V. cholerae* strains minimally encode their T6SS^i1^ in three genetic loci [25,26] (Fig. 1b). The Large cluster encodes the majority of the structural components of the system, including the membrane complex that anchors the system to the bacterial inner membrane *(vasDFK/tssJLM),* the baseplate complex from which the sheath is extended *(hsiF/tssE, vasABE/tssFGK),* the contractile sheath components *(vipAB/tssBC),* and two *tssA-*type proteins involved in the regulation of sheath extension and firing (*vasJL*) [27–29] (Fig. 1a,b). The Large cluster also encodes a VgrG spike *(vgrG-3)* with a specialized bactericidal C-terminus and its cognate immunity factor *(tsiV3)* [12]. Two auxiliary T6SS clusters (Aux1 and Aux2) each encode an Hcp protein *(hcp-1, −2)* that completes the system by forming the central tube upon which the spike is fired out of the cell, an alternate VgrG spike *(vgrG-1, vgrG-2),* a chaperone protein for effector loading *(tap-1, vasW*) [30], and a distinct effector-immunity pair. The VgrG-1 protein encoded by Aux1 is sometimes fused to a specialized C-terminal actin-crosslinking domain (ACD) with anti-eukaryotic properties [4]. Some *V. cholerae* strains carry additional T6SS loci, of which four have been identified [31–34]. Aux3 minimally encodes an effector-immunity pair *(tseH-tsiH)* and a PAAR adaptor necessary for its loading onto the T6SS [35], while Aux4, Aux5, and Aux6 encode their own Hcp, VgrG, and chaperone along with effector-immunity pairs. Aux3 and Aux4 are both carried on mobile genetic elements and are transferred between strains [32,36].

The *V. cholerae* species is a diverse collection of strains that display varying degrees of virulence to humans, including harmless strains, opportunistic pathogens, and pathogenic strains that have evolved to infect the human gastrointestinal tract. Once ingested into the host gut, some pathogenic *V. cholerae* strains cause the deadly secretory diarrhea known as cholera. These toxigenic strains are primarily defined by the presence of the virulence factors cholera toxin and toxin co-regulated pilus [37–40] and have variable serotypes. Most of these strains belong to the pandemic generating lineage, a monophyletic group of strains descended from an ancestor with the O1 serotype [41]. The first six cholera pandemics were caused by O1 strains of the Classical biotype, and the current 7^th^ pandemic is caused by O1 strains of the El Tor biotype [42–45]. T6SS regulation and composition vary between O1 Classical, O1 El Tor, and environmental strains of *V. cholerae.* Most environmental *V. cholerae* strains constitutively express their T6SS [46,47]. For pandemic strains, the O1 El Tor T6SS is tightly regulated by host signals [48], while O1 Classical strains lack a functional T6SS due to mutations in *vipA/tssB, hsiF/tssE, vasE/tssK,* and *vasK/tssM* [9,49].

While the T6SS on a whole is conserved across *V. cholerae*, different strains encode different effectors [24,26]. Environmental *V. cholerae* strains encode a wide variety of effector types, while pandemic *V. cholerae* strains all encode an identical set of distinct effectors referred to as A-type *(tseL, vasX, vgrG-3)* as well as the *vgrG-1* ACD (Fig. 1c). It is important to note that different effector genes at the Aux1 (A or C type) and Aux2 (A-E type) locus encode distinct proteins, while variable types at the Large cluster (A-L) are different C-terminal extensions on a conserved VgrG spike [26]. There is no cross protection between types, and any disagreement in effector set composition can lead to intraspecies competition [26] (Fig. 1a,d). Not only are environmental *V. cholerae* T6SS effector sets highly variable, but T6SS effectors are horizontally transferred [24,50]. One indication of effector transfer between strains is the maintenance of past immunity genes without their cognate effectors (orphan immunity genes) at the *V. cholerae* T6SS loci [24]. The exact mechanism of recombination that leads to orphan retention is unknown, but we hypothesize that orphan retention is an advantageous evolutionary mechanism that allows a constitutive T6SS-producing strain to exchange an effector-immunity type without becoming vulnerable to surrounding ex-kin cells. The Aux1 locus of pandemic *V. cholerae* encodes a single orphan C-type immunity gene, leading us to hypothesize that the Aux1 A-type effector/immunity pair *(tseL/tsiV1)* was acquired by an ancestor of the *V. cholerae* pandemic clade that encoded a C-type effector/immunity pair at Aux1. Acquisition of A-type effectors may have been advantageous for pandemic spread of the clade.

In this study, we aimed to identify the evolutionary history of pandemic *V. cholerae*-associated A-type T6SS effectors. We showed that *V. anguillarum* and closely related fish-colonizing *Vibrio* species ubiquitously encode the Aux1 A-type effector *tseL.* Further, many *V. anguillarum* strains also encode the vgrG-1 ACD, *vasX,* and *vgrG-3.* We indicated likely interaction between *V. cholerae* and *V. anguillarum* in the environment and highlighted potential transfer events of T6SS effectors between the two species. Finally, we showed that the *tseL-tsiV1* effector immunity pair from *V. cholerae* and the Aux1 A-type pair from *V. anguillarum* cross-neutralize each other, indicating an evolutionary relationship between the two clusters. This potential niche sharing and gene flow between *V. cholerae* and *V. anguillarum* may bolster fish as an important environmental reservoir for *V. cholerae*.

## RESULTS

### All *V. cholerae* T6SS loci with the exception of Aux5 and Aux6 are conserved in a clade of fish-colonizing Vibrio species

While the T6SS^i^ is highly conserved across the *Vibrio* genus, different species encode different subtypes of the T6SS^i^. We first aimed to identify *Vibrio* species that encode the core and auxiliary T6SS loci identified in *V. cholerae.* We compiled a dataset of 247 genomes from *V. cholerae (Vch,* n=61), *V. paracholerae (Vpch,* n=12), *V. metoecus (Vmet,* n=26), *V. mimicus (Vmim,* n=22), *V. parilis (Vpar,* n=1), *V. furnissii (Vfur, n=*8), *V. fluvialis (Vflu,* n=18), *V. anguillarum (Vang,* n=63), *V. ordalii (Vord,* n=2), *V. vulnificus (Vvul,* n=9), *V. parahaemolyticus (Vphl,* n=5), *V. scophthalmi (Vsco,* n=4), *V. kanaloae (Vkan,* n=12), and closely related *Vibrio* species *(Vibrio* sp., n=3) and probed for T6SS clusters. Our search identified a single T6SS^i1^ cluster in all species but *Vul.* Strains of *Vphl* (4/5), *V. flu* (2/18), *Vfur* (8/8), *Vang* (39/63), *Vch* (1/61), and *Vmim* (1/22) were also found to encode a T6SS^i5^ (Fig. 2a,b, Additional File 3). All analyzed *Vvul* strains encode a T6SS^i5^ similar to the T6SS^i5^ locus in *Vphl,* while two *Vvul* strains carry an extra T6SS^i5^ with homology to the T6SS^i5^ found in *Vflu, Vfur, Vang, Vch,* and *Vmim* (Fig. 2a, Additional File 3). Based on the reported dispersion of the latter T6SS^i5^ in *Vvul* [51] and our cross-species analysis, this entire locus is likely horizontally transferred.

**Figure 2.**
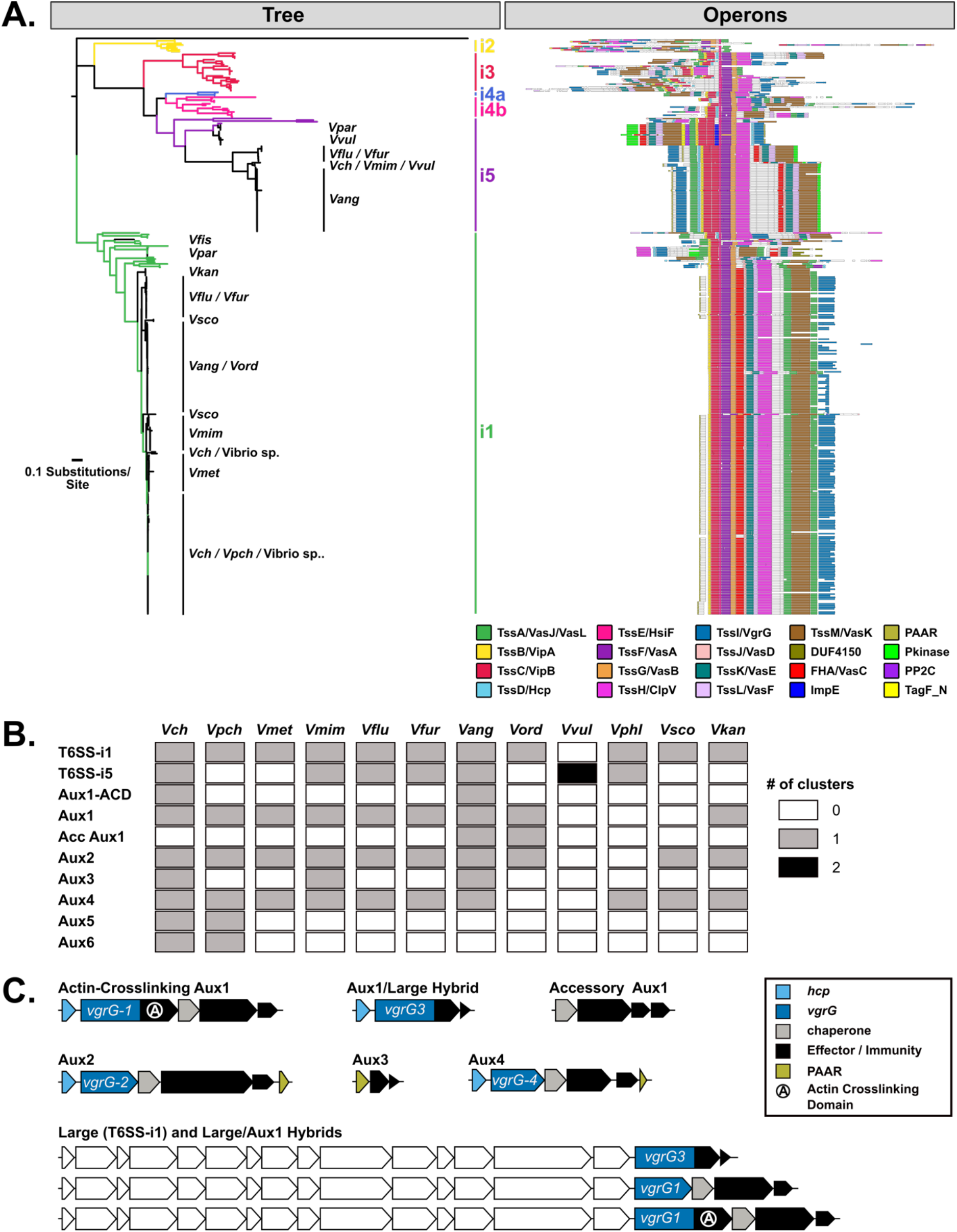
*V. anguillarum* encodes most known *V. cholerae* T6SS loci and a single unique T6SS cluster. **a** Hamburger ((https://github.com/djw533/hamburger) analysis of all analyzed *Vibrio* genomes showing all identified complete T6SS clusters. Grouping of the identified T6SS clusters with known T6SS types is shown by phylogenetic tree (left) and operon structure is indicated (right). Colored branches of the tree indicate Hamburger-internal representative T6SSs corresponding to the indicated type, and black branches indicate input genomes. Operon diagrams are centered around *vipA/vipB.* **b** Presence-absence heatmap of all T6SS clusters identified in *V. cholerae* and *V. anguillarum* and their distribution across all analyzed *Vibrio* species. **c** Schematic diagrams of T6SS clusters identified in *V. anguillarum* showing known *V. cholerae* clusters, *V. anguillarum-specific* hybrid clusters, and a unique *V. anguillarum* locus (Accessory Aux1).

Next, we identified potential auxiliary clusters (containing some but not all elements of a functional T6SS) in each species. To date, six Auxiliary clusters (Aux1-6) have been identified in *Vch* (Fig. 1b). We performed tblastn searches against our *Vibrio* genome dataset for *vgrG* sequences with a 60% grade cutoff (a weighted score accounting for query coverage, e-value, and pairwise identity) as well as sequences for all known effector types from each *Vch* T6SS Aux cluster (30% grade cutoff for Aux effectors and 60% grade cutoff for Large effectors) [24,31–34]. The core *Vch* T6SS loci Aux1 and Aux2 have previously been identified in *Vmet, Vmim, Vflu,* and *Vfur* [23,24]. Our searches confirmed the Aux1 and Aux2 clusters in these species and identified Aux1 and Aux2 loci in *Vang* and *Vord* species (Fig. 2b,c, Additional File 4). In the majority of *Vang* strains, a region homologous to the *Vch* Aux1 cluster from *vgrG-1* through *tsiV-1* is encoded at the end of the Large T6SS^i1^ cluster (Large/Aux1 Hybrid) while a *vgrG-3* homolog is encoded next to the *hcp* gene usually associated with Aux1 (Aux1/Large Hybrid) (Fig. 2c). These hybrid clusters were likely produced by a recombination event between the conserved 5’-regions of *vgrG-1* at the Aux1 cluster and *vgrG-3* at the Large cluster. To our knowledge, this is the first observation of recombination between regions encoding structural components of two T6SS clusters in a *Vibrio* genome. The *Vang* species also encodes Aux3 (2/63) and Aux4 (18/63). Interestingly, *Vang* and *Vord* species encode an extra T6SS cluster absent from all other analyzed species. This novel cluster lacks *hcp* and *vgrG* genes, encoding only a putative DUF4123-containing T6SS chaperone protein [30], a single effector gene and anywhere from 1 to 7 immunity cassettes (Fig. 2b,c). As this cluster always encodes an Aux1 A-type effector homologous to *Vch* TseL (Additional File 1: Figure S2), we have named it Accessory Aux1 (Acc Aux1).

None of the analyzed *Vvul* genomes encode T6SS Aux clusters, and only one of the five analyzed *Vphl* genomes (CFSAN018762) encodes an Aux cluster (Aux4). Three of the four available *Vsco* genomes encode a T6SS, and two of these genomes were found to encode multiple Aux clusters. These two *Vsco* strains were found to encode an Aux2 (2/2) and an Aux4 (1/2) cluster (Additional File 1: Figure S2). The remaining Aux loci in these strains did not encode effectors with *Vch* homologs (data not shown) and were thus exclude from downstream analyses. *Vsco* strain VS-05 was also found to encode Aux4 (Additional File 1: Figure S2), but the Aux4 cluster genes came at the end of an intact T6SS^i1^ cluster carried on a plasmid (data not shown). Five of the 12 analyzed *Vkan* genomes encode a T6SS^i1^, and each of these strains encodes some combination of Aux1, Aux2, and Aux4 (Additional File 1: Figure S2).

### Cross-species effector gene distribution indicates the presence of pandemic-associated *V. cholerae* T6SS effectors in the *V. anguillarum* clade

With our finding that the *Vch* Aux1, Aux2, and Large loci are core T6SS loci in *Vang* and *Vord,* we next aimed to assess the relationship of the *Vang/Vord (Vang* clade) genomes in our dataset to the selected *Vch*, *Vpch, Vmet, Vpar,* and *Vmim (Vch* clade) genomes. We first constructed a multi-species, core genome phylogeny based on 1,889 core proteins (Fig. 3). The *Vang* clade branches from a common ancestor that went on to found the *Vch* and *Vflu/Vfur* clades, indicating that *Vang* and *Vord* are more distant relatives of *Vch* than are *Vflu* and *Vfur.* We next assessed the distribution of known *Vch* effector types across the selected *Vibrio* species. We identified *Vch* effector types in the target genomes by tblastn with known representatives for each effector type using a 30% grade cutoff for Aux cluster effectors and a 60% grade cutoff for VgrG effectors. Individual strain effector sets are shown in the expanded phylogenetic trees for each species in the supplement (Additional File 1: Figure S1, S2). The distribution of effector types at the Aux1 cluster (Fig. 3) indicates that the C effector-immunity pair is widespread across the *Vch*, *Vflu/Vfur,* and *Vang* clades and was likely encoded by the founding Aux1 cluster in the most recent common ancestor of the three clades. The Aux1 A effector (*tseL*), however, is irregularly distributed across the three clades, indicating likely horizontal transfer of this effector-immunity module (Fig. 3). Previous work has demonstrated that the Aux1 A effector is highly enriched in the pandemic clade of *Vch* and is sporadically distributed throughout environmental *Vch*, *Vmet,* and *Vmim* strains [24]. Here we show for the first time that the Aux1 A effector is widespread within the *Vang* clade. Interestingly, *Vang* and *Vord* uniformly encode both and Aux1 A and C effectors. This excess of Aux1 effectors is due to the presence of both a standard Aux1 cluster and the newly identified Acc Aux1 cluster (Fig. 2b,c).

**Figure 3.**
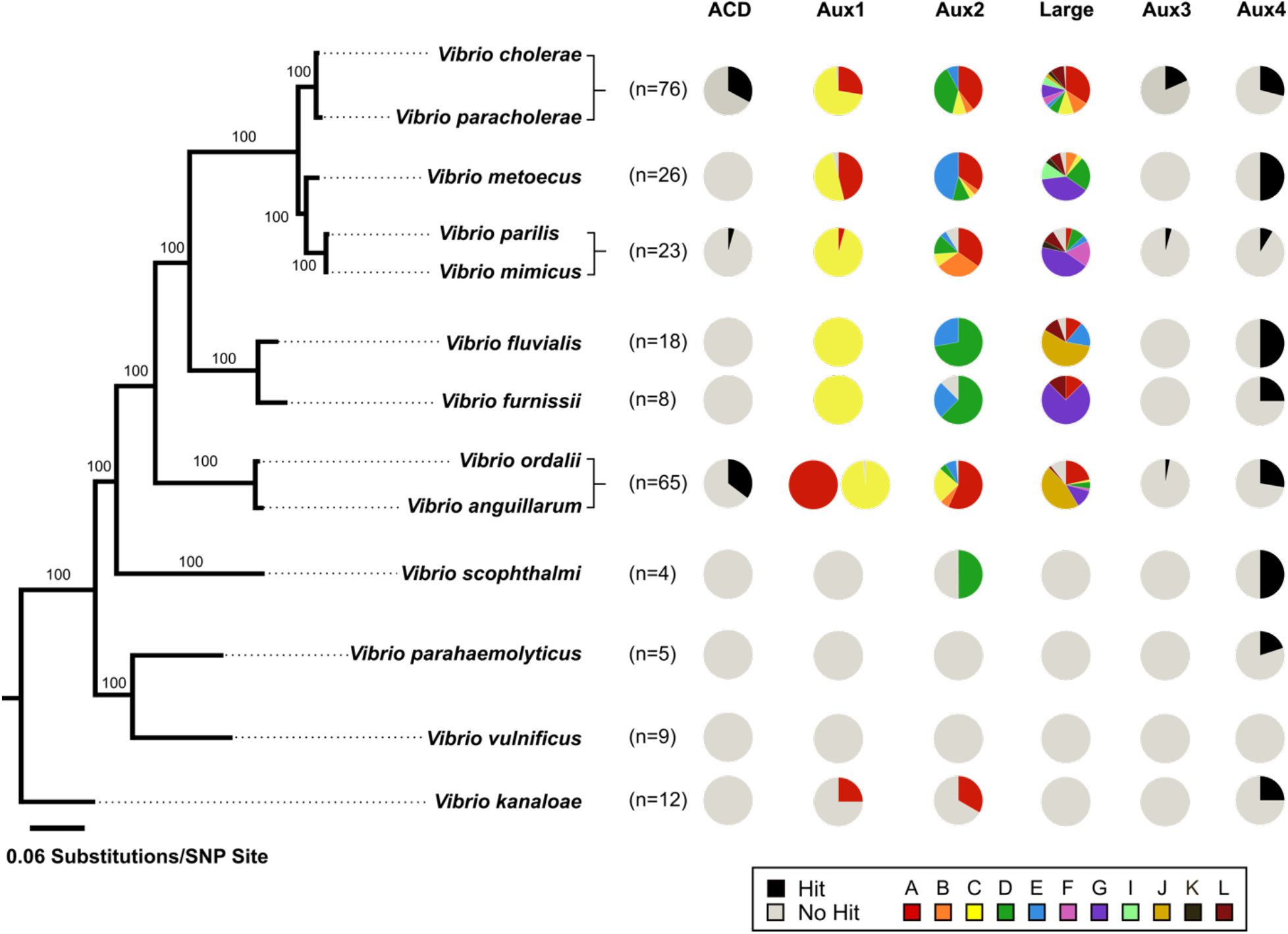
Known *V. cholerae* T6SS effector types, including pandemic-associated effectors, are found in *V. anguillarum.* (Left) Maximum-likelihood tree built on 1,889 core proteins from 30 *Vibrio* genomes collapsed by species. Branch support was calculated using 100 bootstrap replicates. Support values are indicated. (Right) Pie charts indicating the presence of each known *V. cholerae* Aux1, Aux2, and Large effector type (Fig. 1c) as well as the actin-crosslinking domain of *vgrG-1,* the Aux3 effector *tseH* and the Aux4 effector *tpeV.* Number of genomes included for each species (n) is indicated.

A similar distribution is seen for other pandemic-associated *Vch* effectors. The *vgrG-1* ACD is only observed in *Vch*, *Vpar*, and the *Vang* clade: 25 of 61 *Vch* strains, 1 of 1 *Vpar* strains (*V. parilis* RC586) [52], and 23 of 63 *Vang* strains (Fig. 3, Additional File 1: Figure S1, S2). At the Aux2 cluster, D- and E-type effectors are present in all three clades, but A- *(vasX),* B-, and C-type effectors are present only in the *Vch* and *Vang* clades (Fig. 3). The A-type effector encoded at the large cluster *(vgrG-3),* however, is found in the *Vflu/Vfur* clade as well as the *Vch* and *Vang* clades (Fig. 3). Finally, the Aux3 effector/immunity pair *(tseH/tsiH),* which is enriched in pandemic *V. cholerae* strains [36], is restricted to *Vch*, *Vmim*, and the *Vang* clade. It is important to note, that our analysis is restricted by the small number of available *Vflu* and *Vfur* genomes. It is possible that the observed lack of pandemic-associated *Vch* effectors in the *Vflu/Vfur* clade is due to sampling bias. Due to the presence of the Aux1 A effector (*tseL*) at a novel T6SS gene cluster that is widespread in the *Vang* clade, we focused on the evolutionary history of this effector gene for the remainder of the study.

### Aux1 A-type T6SS effectors are horizontally transferred between *V. anguillarum* and *V. cholerae*

We next sought to identify horizontal gene transfer events between *Vch* and the *Vang* clade. Every identified A-type Aux1 cluster in *Vch*, *Vpch, Vmet,* and *Vmim* also carries at least one C-type orphan immunity gene, indicating that these strains are derived from one or more Aux1 C ancestors and that the Aux1 A effector was acquired from elsewhere. The *Vang* group is of particular interest as an Aux1 A donor due to the presence of the Acc Aux1 cluster, which does not encode any C-type orphans and was therefore likely founded with an A-type effector.

To identify potential donor Acc Aux1 clusters, we extracted all identified A-type effector and immunity genes from each species and clustered the corresponding amino acid sequences with a cut-off of 80%. This analysis indicated that the Aux1 A effector can be grouped into five subtypes (A1-A5) (Additional File 1: Figure S3). Aux1 A2 and A3 are unique to the *Vang* clade, while A1 and A4 are shared between *Vch* and the *Vang* clade. The Aux1 A immunity genes are more diverse than their cognate effectors (Additional File 1: Figure S4). To account for this increased amino acid diversity, we lowered our clustering cut-off to 70%. The resulting clusters lead us to further subdivide Aux1 A1 and A2 subtypes into A1a/b and A2a/b, respectively. We next built a single protein phylogeny of the Aux1 A effectors and overlaid the corresponding Aux1 and Acc Aux1 clusters for each strain or group of strains along with their new effector-immunity subtypes (Fig. 4a). From this tree, it is evident that the Aux1 A1a effector-immunity pair found in the *Vch* and *Vpch* strains most likely originated from a group of *Vang* strains.

**Figure 4.**
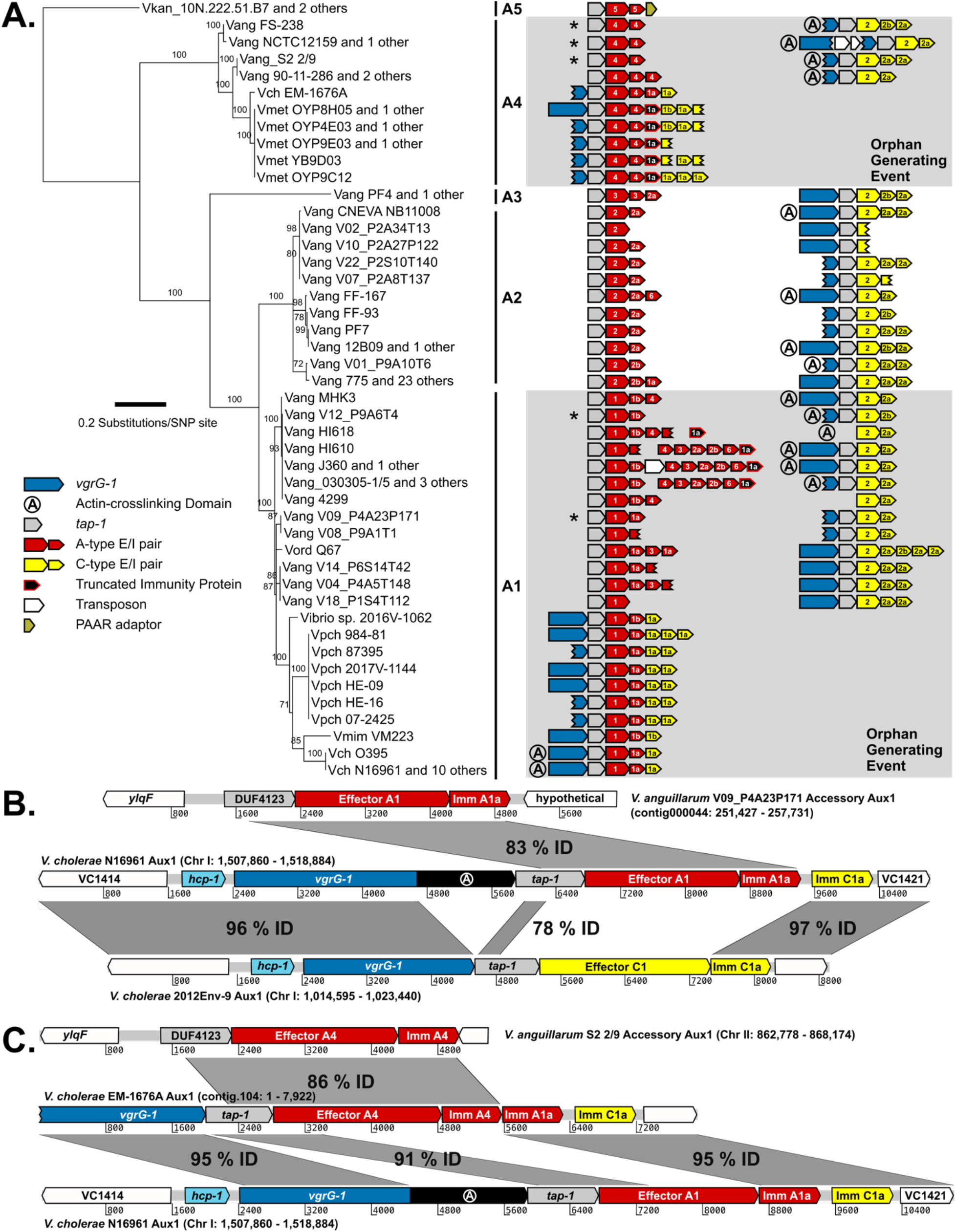
*V. anguillarum* Accessory Aux1 was horizontally transferred into *V. cholerae* recipient strains. **a** Maximum-likelihood tree of identified Aux1 A-type effectors (TseL). Bootstrapping support values for each node are shown. Accessory Aux1 and/or Aux1 schematic for each strain is shown on the right. Effector/immunity type is indicated by color, and each effector/immunity cassette is labeled with its associated subtype. Grey boxes indicate groups of strains potentially involved in transfer of the A-type effector from *V. anguillarum* and the generation of an orphan immunity gene in *V. cholerae.* Asterisks mark the *V. anguillarum* Accessory Aux1 clusters most likely to have been transferred to *V. cholerae.* **b** Artemis alignment of the T6SS Aux1 cluster from pandemic *V. cholerae* strain N16961 with its potential precursor loci – Accessory Aux1 from *V. anguillarum* V09_P4A23P171 and Aux1 from *V. cholerae* 2012Env-9. This alignment represents the potential orphan generating transfer event indicated in (a, bottom grey box). **c** Artemis alignment of the T6SS Aux1 cluster from *V. cholerae* strain EM-1676A with its potential precursor loci – Accessory Aux1 from *V. anguillarum* S2 2/9 and Aux1 from *V. cholerae* N16961. This alignment represents the potential orphan generating transfer event indicated in (a, top grey box). For Artemis alignments, regions of homology are indicated by grey boxes, which are labeled with the nucleotide identity between the covered regions.

Alternatively, it is possible that *Vpch* acquired the A1a effector-immunity pair from *Vang* and subsequently transferred the genes to *Vch*. A group of *Vang* and *Vord* strains *(Vang* V09_P4A23P171, *Vang* V04_P4A5T148, *Vang* V14_P6S14T42, and *Vord* Q67) encode A1a immunity genes ranging from 89.8 to 91.1% nucleotide identity and 91.7 to 92.5% amino acid identity to the pandemic *Vch* A1a immunity gene (Additional File 1: Figure S4). All of the *Vpch* strains encoding an Aux1 A1a effector-immunity pair, except *Vpch* 877-163, encode an A1a immunity gene with 91.7% nucleotide identity and 89.5% amino acid identity to the pandemic *Vch* A1a immunity gene (Additional File 1: Figure S4). *Vpch* 877-163 encodes an Aux1 A1a effector-immunity pair nearly identical to the pandemic *V. cholerae* Aux1 locus, but we cannot determine the directionality of transfer (Additional File 1: Figure S4). Based on A1 a immunity gene homology, we are unable to decern which species was the terminal donor of the Aux1 A1 effector to *Vch*, but our data support *Vang* as the initial donor to the *Vch* clade.

All *Vmet* strains with an Aux1 A effector as well as *Vch* EM-1676A encode an A4 effector-immunity pair, which most likely originated from a different sub-clade of *Vang* strains including *Vang* S2 2/9, *Vang* NCTC12159, and *Vang* FS-238 (Fig. 4a). The Aux1 A1b effector found in the *Vch*-related Vibrio sp. 2016V-1062 and *Vmim* VM223 may have originated from a group of several A1b-encoding *Vang* strains, including *Vang* V12_P9A6T4 (Fig. 4a).

### Recombination between Aux1 and Accessory Aux1 to generate orphan immunity arrays occurred on at least two separate instances

We aimed to use the Aux1 C-type orphan immunity genes present in the *Vch* clade to identify groups of strains that constitute likely recipients of the Aux1 A effector-immunity module from the *Vang* clade. To identify potential Aux1 recipient clusters, we extracted all identified C-type effector and immunity genes, including both bona fide (encoded next to their cognate effector) and orphan immunity genes, from each species and clustered the corresponding amino acid sequences with a cut-off of 80%. Like the A-type effectors, we identified five C effector subtypes (C1-C5) (Additional File 1: Figure S5). Again, the cognate immunity genes for two C subtypes were more variable than their effectors, leading to further subdivisions (C1a/b and C2a/b) (Additional File 1: Figure S6). Unlike the Aux1 A effectors, C effector subtypes are clade specific, with C1 restricted to the *Vch* clade, C2 restricted to the *Vang* clade, and C3-5 restricted to the *Vflu/Vfur* clade.

We next overlaid our constructed Aux1 A effector phylogeny with our newly established Aux1 C effector-immunity sub-types (Fig. 4a). Each orphan-encoding Aux1 A cluster in *Vch*, *Vpch, Vmet,* or *Vmim* carries a C1 type orphan. Based on this result, we concluded that on one or more occasions the Acc Aux1 cluster from a *Vang* strain was transferred to *Vch* and closely related *Vibrio* species where it recombined with an Aux1 C1 cluster (Fig. 4b,c, Additional File 1: Figure S7). An alternate scenario in which Acc Aux1 recombined with the Aux1 C2 cluster in the *Vang* clade to create the orphan-encoding locus we see in *Vch* is unlikely based on the orphan C immunity subtypes.

The *Vch* 2012Env-9 Aux1 cluster is a potential predecessor of the pandemic Aux1 locus, as it encodes a bona fide C immunity protein that is >95% identical to the orphan C immunity protein encoded by pandemic strains (Fig. 4b). *Vch* 2012Env-9 is also a member of a sister clade to the *Vch* pandemic clade (Fig. 1c) and is thus a good extant relative of the pre-pandemic ancestor that received the Aux1 A effector [24]. Recombination between Acc Aux1 and Aux1 likely occurred by homology-facilitated illegitimate recombination, a known mechanism for interspecies gene transfer [52–55]. An initial crossover occurred in a region of strong homology between the Acc Aux1 DUF4123-containing chaperone and the Aux1 *tap-1* chaperone. The second crossover event likely occurred between the 3’ end of the donor A1 immunity gene and the 3’ end of the C1 effector, despite no obvious homology between the two genes, leading to displacement of the C1 effector and maintenance of the C1 orphan (Fig. 4b).

All Aux1 A4-encoding *Vmet* strains and *Vch* EM-1676A encode not only a C1 orphan, but also an A1 type orphan (Fig. 4a) [24]. This array of orphan immunity genes indicates that transfer of the Acc Aux1 A4 effector/immunity pair into *Vmet/Vch* was likely a second independent transfer event in which the incoming Acc Aux1 A4 cluster recombined into an Aux1 cluster that had already acquired the Aux1 A1 effector/immunity pair (Fig. 4c). There is no obvious recipient Aux1 cluster into which Acc Aux1 A4 recombined to form the *Vch* EM-1676A Aux1. The *Vch* EM-1676A Aux1 cluster encodes type C1a and A1a orphan immunity genes (Fig. 4a,c) but no *vgrG-1* ACD. Based on this, we hypothesize that Acc Aux1 A4 either recombined into an unidentified Aux1 cluster with a matching immunity gene array but no ACD or a pandemic-like Aux1 that later lost its ACD. The necessary gene clusters to back either hypothesis, while likely present in the *V. cholerae* species, are yet to be discovered.

### *V. cholerae* kills *V. anguillarum* in a T6SS-dependent manner

We hypothesize that *Vch* and *Vang* co-occupy a niche in the aquatic reservoir and compete in a T6SS-dependent manner, leading to the subsequent exchange of genetic material. To test this hypothesis, we assessed whether *Vch* can kill *Vang* via its T6SS by competing two strains of *Vch* with constitutively active T6SSs, the O37 pathogenic strain V52 and the environmental strain DL4211, against a *Vang* isolate (VIB43). The T6SS effector set of *Vang* VIB43 (Aux1 C2, Acc Aux1 A1, Aux2 C, Large G) is incompatible with both V52 (Aux1 A1, Aux2 A, Large A) and DL4211 (Aux1 C1, Aux2 E, Large E). After 4 hours of co-incubation at 28°C on agar at a 1:1 ratio, both strains of *Vch* outcompeted *Vang* VIB43 (Fig. 5a-d). Importantly, co-incubation of *Vang* VIB43 with *Vch* carrying an in-frame deletion in the T6SS membrane complex component *vasK* (V52 Δ*vasK*and DL4211 Δ*vasK*) shows no competitive advantage for either strain (Fig. 5a-d), indicating that this fitness advantage for *Vch* is T6SS-dependent. In theory, *Vang* VIB43 should also kill both *Vch* strains, but our understanding of the T6SS induction conditions for *Vang* is limited. It is possible that the *Vang* VIB43 T6SS was not induced under our experimental conditions.

**Figure 5.**
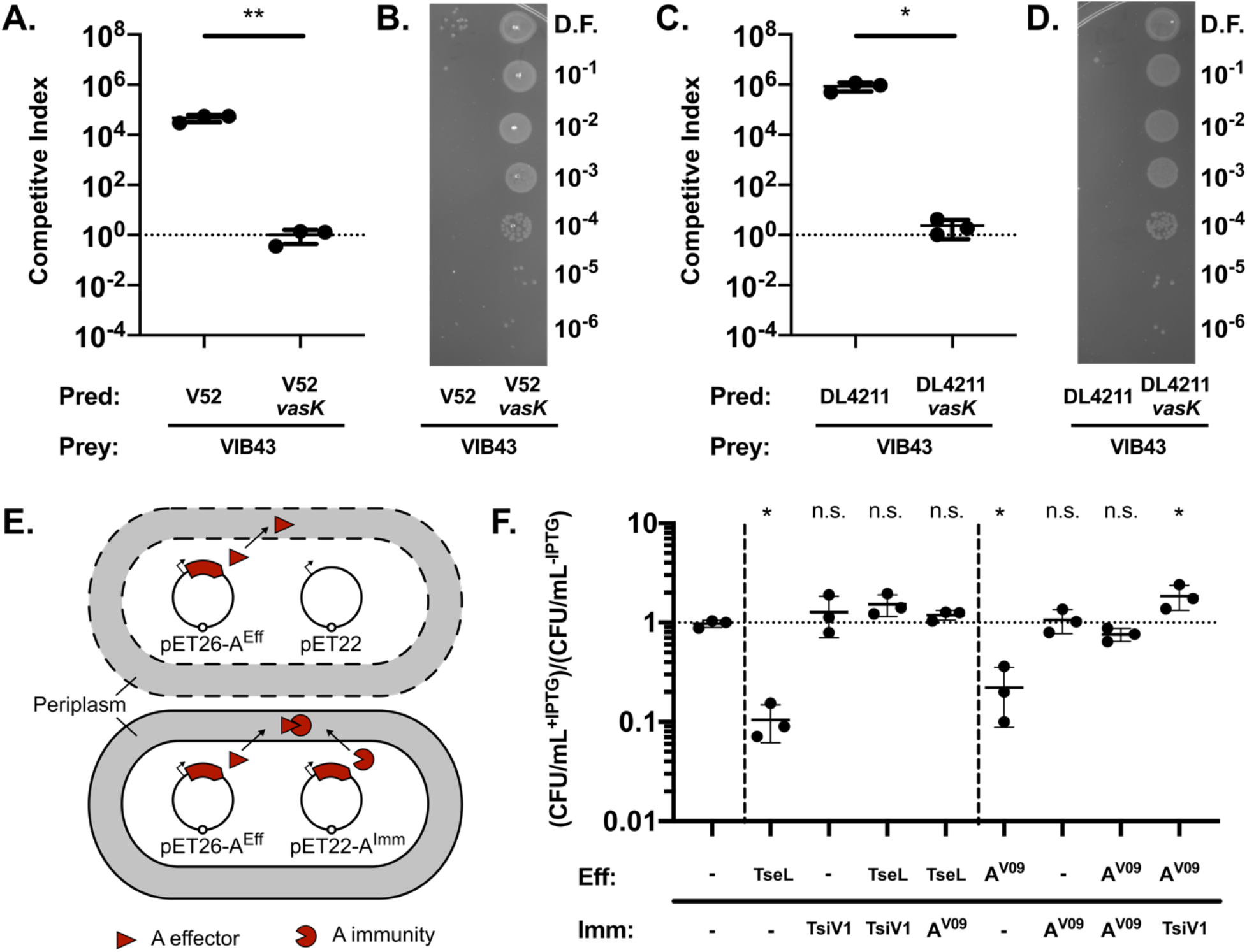
A-type effector/immunity pairs from *V. cholerae* Aux1 and *V. anguillarum* Accessory Aux1 are cross-neutralizing. **a** Competitive killing assay results for *V. cholerae* strain V52 (T6SS on) or V52 *DvasK* (T6SS off) vs *V. anguillarum* strain VIB43. **b** Representative image of dilution spots on agar plates selecting for VIB43 prey cells from competition assays against V52 or V52 *DvasK.* **c** Competitive killing assay results for *V. cholerae* strain DL4211 (T6SS on) or DL4211 *DvasK* (T6SS off) vs *V. anguillarum* strain VIB43. **d** Representative image of dilution spots on agar plates selecting for VIB43 prey cells from competition assays against DL4211 or DL4211 *DvasK.* **a,c** Significance was determined by unpaired t test (* p = 0.0115, ** p = 0.0056). **e** Diagram of *E. coli* BL21(DE3) dual-expression viability assay. Dashed cell membranes represent lysis. **f** Dual-expression viability assay with results plotted as the ratio of CFU/mL recovered with induction (+IPTG) to CFU/mL recovered without induction (-IPTG). TseL = *V. cholerae* A1 effector. TsiV1 = *V. cholerae* A1a immunity protein. A^V09^ = *V. anguillarum* A1 effector or A1a immunity protein. Significance was determined by one-way ANOVA with Dunnet’s multiple comparisons test (* p = (0.0190, 0.0486, 0.0182), n.s. = non-significant). All comparisons were made to strain carrying two empty vectors. **a,c,f** Quantitative results are from three independent experiments (n=3). Individual replicates are shown. Horizontal bars represent the mean, and error bars represent SD.

### *V. cholerae* Aux1 A1a and *V. anguillarum* Acc Aux1 A1a effector/immunity pairs crossneutralize

If the *Vch* Aux1 A1a effector-immunity pair originated from the *Vang* Acc Aux1 gene cluster, then each immunity gene should be cross-protective against the effector of the opposing cluster. To test this hypothesis, we co-expressed either the A-type effector from *Vch* N16961 (TseL) or *Vang* V09_P4A23P171 (Aeff^V09^), each fused to an N-terminal periplasmic secretion signal, with both their cognate immunity gene and the opposing immunity gene (TsiV1 or Aimm^V09^ respectively) in *E. coli* BL21(DE3) (Fig. 5e). Expressing either TseL or Aeff^V09^ alone leads to an approximately 10-fold reduction in the number of recovered, viable *E. coli* cells, while expression of either TsiV1 or Aimm^V09^ alone had no effect on cell viability (Fig. 5f). Cells co-expressing TseL or Aeff^V09^ with their respective cognate immunity genes had no significant reduction in viability. Neither was a reduction in viability observed when each effector was co-expressed with the opposing immunity gene. These results support our hypothesis that TseL and Aeff^V09^ are related effector-immumity clusters that can cross-neutralize each other.

## DISCUSSION

### Inferring the interaction of *V. cholerae* and *V. anguillarum* in nature

*V. cholerae* and *V. anguillarum* are both aquatic organisms that undergo a pathogenic cycle, and it is possible that these two species share common niches. Previous studies have demonstrated, both *in-situ* and *in-silico,* that *V. anguillarum* and *V. cholerae* come into contact in the environment. It has been shown experimentally that *V. cholerae* can maintain R-plasmids received via conjugative transfer from *V. anguillarum* [56], indicating that *V. cholerae* and *V. anguillarum* have the potential to share DNA. While R-plasmids have a broad host range, it is interesting to note that *V. cholerae* cannot stably maintain R-plasmids received from *Enterobacteraceae* [56]. More recently, it was discovered that a genomic island carried in four strains of O1 El Tor *V. cholerae* is 97% identical at the nucleotide level to a homolog in *V. anguillarum* VIB43 [57]. Similar loci have also been identified in multiple strains of non-O1 *V. cholerae* [58]. Our findings further support the idea that *V. cholerae* interacts with *V. anguillarum,* as we identify specific gene transfer events between the two species. We also show that should *V. cholerae* and *V. anguillarum* come into contact in the environment, *V. cholerae* can efficiently kill *V. anguillarum* with its T6SS. The DNA released from lysed *V. anguillarum* cells can then be taken up by *V. cholerae* [59]. This is likely not a one-way interaction, as we show that *V. anguillarum* encodes its own arsenal of T6SS effectors. The *V. anguillarum* T6SS does not appear to be active in strain VIB43 under our experimental conditions, but the *V. anguillarum* T6SSs have been shown to be differentially active under conditions mimicking marine or intra-host conditions and are effective at killing prokaryotic competitors [22,60].

*A* niche of interest for the interaction of *V. cholerae* and *V. anguillarum* is the gut and skin of fish. While *V. anguillarum* is primarily known as a fish pathogen, the incidence of *V. cholerae* colonizing fish in the wild is less understood. It has been shown, however, that both *V. anguillarum* and *V. cholerae* are highly chemotactic towards rainbow trout intestinal mucus [61]. It is possible that the fish gut provides a stable niche for interaction, interspecies killing and gene transfer. Of the 23 known *V. anguillarum* serotypes, only O1, O2, and O3 cause vibriosis in fish [62]. The remaining serotypes are comprised of mostly non-pathogenic environmental isolates from sediment, phytoplankton, and zooplankton. The chitinous exoskeleton of zooplankton is also a niche of interest for the interaction of *V. cholerae* and *V. anguillarum,* as chitin metabolism by *V. cholerae* induces both the T6SS and the natural competence machinery [59,63]. Less is known about natural competence in *V. anguillarum*, but isolates of this species have been shown to encode chitinase and can grow with chitin and N-acetylglucosamine as their sole carbon source [64]. Cocolonization of zooplankton by *V. cholerae* and *V. anguillarum* could result in bi-directional killing and genetic exchange. Our results do not favor either niche as the site of interaction and gene transfer between these two species. We have shown here, however, that a pre-pandemic *V. cholerae* strain carrying an Aux1 C1 effector potentially acquired the Aux1 A1 effector from *V. anguillarum,* likely as a result of physical contact in the past.

Our results as well as previous data from Kirchberger *et al.,* 2017 [24] show that *tseL* is found in a small number of environmental *V. cholerae* strains and several *V. paracholerae* strains. Approximately half of the analyzed *V. paracholerae* strains encode an A1 effector protein that is closely related to the A1 effector found in pandemic *V. cholerae* (Fig. 4a, Additional File 1: Figure S1). As *V. cholerae* and *V. paracholerae* strains have been isolated from common water sources, an alternate hypothesis to direct interaction between pre-pandemic *V. cholerae* and *V. anguillarum* is gene transfer between *V. anguillarum* and *V. paracholerae* and subsequent transfer to the *V. cholerae* pre-pandemic ancestor. Regardless of which species constitutes the terminal *tseL* donor, our orphan C immunity data supports *V. anguillarum* as the origin of the A-type effector-immunity pair in *Vch* and *Vpch.*

### Potential of the fish as a niche for pandemic *V. cholerae*

*V. cholerae,* including some toxigenic O1 strains, has been isolated from 30 different species of fish [65]. Fish colonization is likely to provide *V. cholerae* with benefits, such as protection from the stresses of the aquatic environment and dissemination across long distances both by the fish harboring the bacteria and the birds that eat the fish [66,67]. In the laboratory, the zebrafish gut is a hospitable environment for *V. cholerae,* as the bacterium readily colonizes the intestine, attaches to the epithelium, and forms microcolonies [68,69]. O1 El Tor infection of the zebrafish leads to a sustained colonization, while infection with O1 Classical strains or non-O1/non-O139 environmental strains leads to transient colonization [68,69]. Further, expression of major human virulence factors CT and TCP appear to be dispensable for zebrafish gut colonization [68]. The T6SS could play a role in colonization, as O1 Classical strains of *V. cholerae* carry an inactive T6SS [9,49] and O1 El Tor strains should express their pathoadaptive T6SS in the fish gut. Interestingly, infection with *V. cholerae* strain AM-19226 (non-O1/non-O139), which constitutively expresses its T6SS, shows a similar transient colonization pattern to O1 classical strains [69], indicating that there is more to fish colonization than simply having an active T6SS. A single study has directly investigated the *V. cholerae* T6SS in the fish gut. In this work, an O1 El Tor strain that was modified to constitutively express the T6SS was shown to displace the resident fish gut microbes in a *vgrG-1* ACD dependent manner [70]. This is not sufficient to explain colonization of the fish gut by O1 El Tor strains, however, because environmental strain AM-19226 also constitutively expresses its T6SS and encodes the *vgrG-1* ACD [24]. We believe that our results warrant investigation of the role of different T6SS effector types, specifically pandemic-associated T6SS effectors like TseL, in fish colonization by *V. cholerae.* Improved colonization of the fish gut could lead to increased dissemination and potentially adaptation to a host intestinal tract.

### T6SS effector gene evolution by interspecies transfer and its role in pandemicity

We show that *V. cholerae* is sharing genetic material, specifically pandemic-associated T6SS effectors with *V. anguillarum* and *V. ordalii*. Common T6SS effectors is likely an indication of a shared niche because acquiring effector/immunity sets similar to neighboring cells is a protective strategy [50]. Here we show that, within our dataset, the *V. cholerae* T6SS Aux1 A-type effector *tseL* is uniquely shared between the *V. cholerae* and *V. anguillarum* clades. It is important to note that *V. anguillarum* is more extensively sampled than *V. fluvialis/V. furnissii,* so we cannot rule out sampling bias in the observed distribution of TseL. Our data show that, on at least two occasions, A-type effectors originating in the *V. anguillarum* Acc Aux1 cluster were transferred to the *V. cholerae* clade (Fig. 6). In the first transfer event, an A1a effector-immunity module was likely acquired by the *V. cholerae* pre-pandemic ancestor, displacing its C1 effector gene (Fig. 6a). In the second transfer event, an A4 effector-immunity module displaced the A1 effector (Fig. 6a). Based on the diversity of orphan C immunity arrays observed at Aux1 in *V. cholerae, V. paracholerae, V. metoecus,* and *V. mimicus,* it is likely that more than two transfers of the Aux1 A effector occurred between the two clades (Fig. 6b).

**Figure 6.**
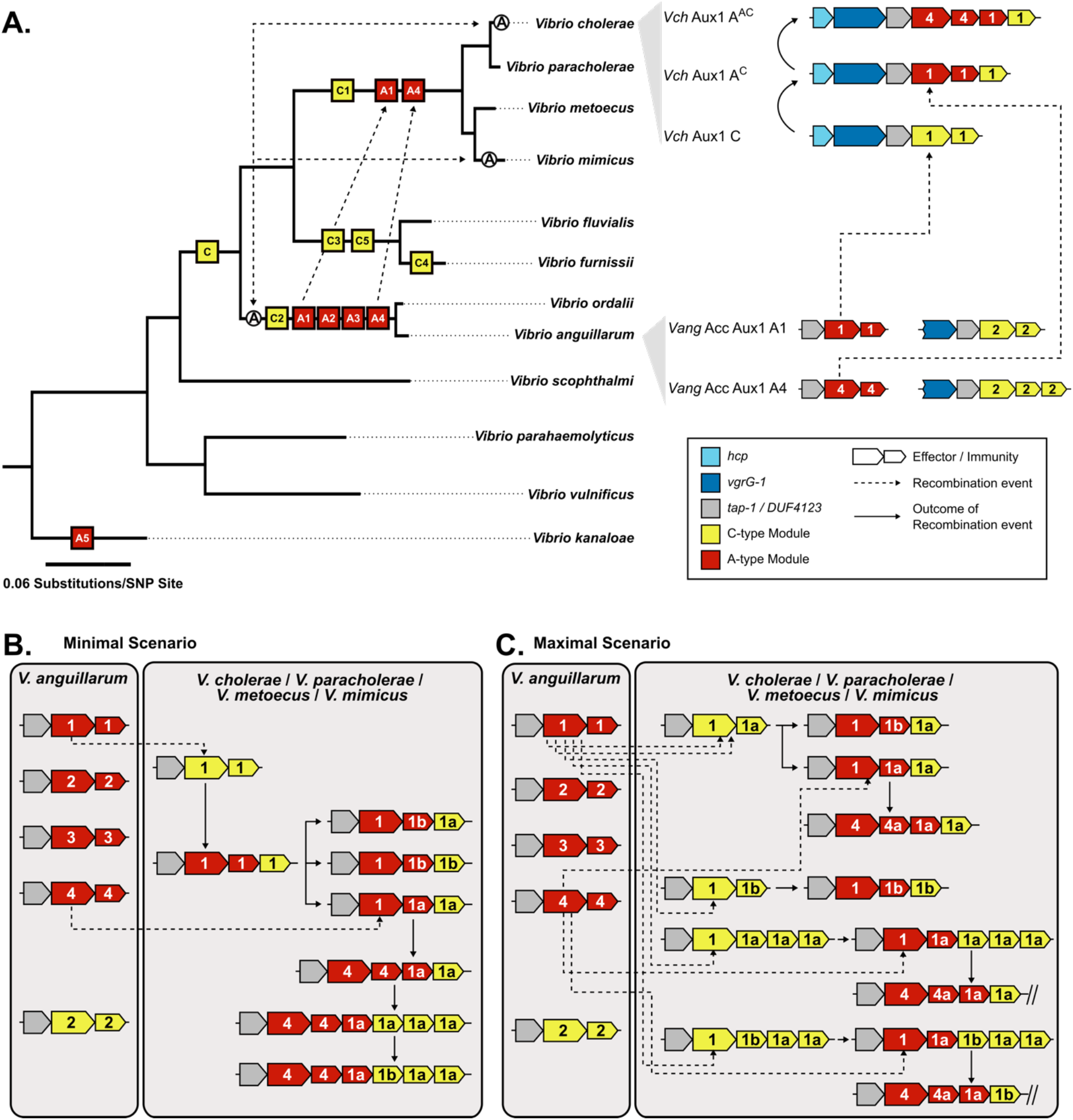
Model of pandemic-associated T6SS effector evolution at the Aux1 cluster of *V. cholerae.* **a** (left) Core genome SNP-based maximum likelihood tree from (Fig. 3) overlaid with likely points of introduction for each Aux1 effector type/subtype and the ACD of *vgrG-1.* Potential horizontal transfer events are indicated by uni- or bi-directional dashed arrows. (right) Representative genetic diagrams of Aux1 clusters in *V. cholerae* as well as Accessory Aux1 and Aux1 clusters in *V. anguillarum.* Effector/immunity types are indicated by color and effector/immunity subtypes are indicated. **b,c** Schematic diagrams representing the potential transfer events leading to the array of Aux1 clusters observed in *V. cholerae, V. paracholerae, V. metoecus,* and *V. mimicus.* **b** Minimal gene transfer scenario consisting of Aux1 A1 transfer, divergence of the recipient cluster, subsequent transfer of Aux1 A4, and further divergence of that cluster. **c** Maximal gene transfer scenario consisting of multiple individual transfer events for both Aux1 A1 and Aux1 A4. **a-c** Effector/immunity types are indicated by color and effector/immunity subtypes are indicated. Dashed arrows indicate recombination events, and solid arrows indicate the resulting cluster from said recombination events.

If *V. cholerae* shares niches and DNA with both *V. fluvialis/V. furnissii* and *V. anguillarum* but only exchanges the Aux1 A T6SS effector with *V. anguillarum,* then it is possible that this effector is adaptive to the shared niche of *V. cholerae* and *V. anguillarum.* This raises the question of whether TseL plays an environmental role, as pandemic-associated *V. cholerae* genomic islands have been shown to have environmental significance [71]. One potential environmental role of TseL could be fish gut colonization, which in turn could favor pandemicity in some way. For instance, the VgrG-1 ACD has been shown to induce intestinal inflammation in the mouse model of infection [8,72] and expedite niche clearance in the fish gut through the induction of peristalsis [70]. This is a clear example of a T6SS effector that is beneficial for both fish gut colonization and mammalian disease. Based on our *in-silico* results showing that TseL is widespread in the *V. anguillarum* clade, we hypothesize that a parallel scenario may exist for the Aux1 A effector. We cannot, however, restrict the potential benefits of TseL to fish gut colonization, as TseL is present in both vibriosis-causing and non-pathogenic, environmental *V. anguillarum* isolates.

## CONCLUSION

In this study, we assessed the distribution of T6SS loci and effector types across 13 *Vibrio* species to reconstruct the evolutionary history of the pandemic *V. cholerae* T6SS and to indicate potential interspecies interactions. Our genomic results indicated sharing of T6SS effector/immunity modules between *V. anguillarum* and *V. cholerae* and lead us to conclude that these two species may share one or more aquatic niches and regularly compete. We propose that *V. anguillarum* constitutes an environmental reservoir of pandemic-associated *V. cholerae* T6SS effectors, indicating that the pandemic A-type effectors may play important roles outside of human pathogenesis. Our findings highlight fish colonization as a potential stage during the evolution of pandemic *V. cholerae* and indicate that competitive fitness in the fish gut may be important for pandemic *V. cholerae* strains in the environmental reservoir.

## METHODS

### Acquisition and annotation of publicly available genome sequences

All genome sequences used in this study are publicly available and were downloaded from the NCBI GenBank database. GenBank Accession numbers for all strains are listed in Additional File 2: Table S1. A single fasta file was generated for each genome and all files were re-annotated with Prokka (v1.12) [73] for uniformity across genomes.

### Identification of T6SS loci and T6SS cluster typing

Complete T6SS clusters were identified using HMmer Based UndeRstandinG of gene clustERs (hamburger)(https://github.com/djw533/hamburger). GFF files generated with Prokka were analyzed with hamburger using the -t option for an automatic T6SS search based on 13 highly conserved structural and regulatory genes. Hamburger then aligned the identified T6SS clusters with an internal reference set of identified T6SS cluster types (T6SS^i1^ – T6SS^i5^) to assign types to the extracted loci. This alignment was then used by hamburger to generate a phylogenetic tree of T6SS loci with corresponding gene diagrams aligned around the *vipA/vipB (tssB/tssC)* cassettes.

### Identification of Auxiliary T6SS clusters

Auxiliary T6SS clusters were not identified by hamburger as they do not encode a sufficient number of T6SS genes to be considered a contiguous T6SS cluster and subsequently extracted by the program using the -t option. Hamburger does not identify effector and immunity gene cassettes as these are extremely variable both between and within species. For these reasons, auxiliary clusters were identified manually from our custom database of 247 genomes. Further, several genomes in this study have only been assembled into contigs, occasionally resulting in fragmented T6SS loci. As there is no one method that will capture every loci under these conditions, we use the following three methods. *Vibrio* auxiliary clusters commonly encode *vgrG* genes, and so auxiliary clusters were identified in an unbiased manner by performing a tblastn search for *vgrG-2* (VCA0018) from *V. cholerae* N16961 with a Geneious grade (a weighted metric combining query coverage (0.50), e-value (0.25), and pairwise identity (0.25)) cutoff of 60%. Auxiliary clusters were also identified in a biased manner by performing a tblastn search for all known T6SS effector types from *V. cholerae* with a Geneious grade cutoff of 30% for effectors associated with Auxilliary clusters and a Geneious grade cutoff of 60% for the VgrG effectors of the Large cluster [24,31–34]. The N-terminal VgrG portion of all *V. cholerae* Large cluster effector types is highly conserved, and thus these effector types require a higher grade cutoff to differentiate. Finally, once auxiliary loci had been identified in each species, flanking, non-T6SS genes were used to find any missing loci in other strains of that species. The identified auxiliary loci were verified against the literature to separate known and novel T6SS loci [23,24]. All identified loci were extracted from their respective genomes for downstream analyses. As this study focuses on *V. anguillarum,* the genomic locus of all identified T6SS auxiliary clusters from *V. anguillarum* strains are listed in Additional File 4.

### Core genome and single protein phylogeny

A species level phylogeny (Fig. 3, Fig. 6a) was constructed as follows: The core genome of 30 *Vibrio* strains (Additional File 2: Table S2) corresponding to 13 described species, including *V. paracholerae,* an emerging new species/subspecies closely related to *V. cholerae,* was extracted using usearch with an amino acid cutoff of 30% and then aligned using MUSCLE [74,75], as implemented by the BPGA pipeline [76]. The resulting alignment of 1,889 core proteins, corresponding to 650,474 amino acid positions, was then used to construct a phylogenetic tree using the GAMMA+WAG substitution model in RAxML (v8.0.26) [77]. Branch support was calculated using 100 bootstrap replicates. Phylogenetic tree was visualized from RAxML-generated newick file using TreeGraph 2 (v2.15.0-887 beta) [78]. Branches with bootstrapping support values <70 were collapsed.

Core genome based phylogenetic trees for individual clades and species (Additional File 1: Figure S1, S2) were constructed as follows: Annotated GFF3 files generated by Prokka (v1.12) [73] were used for this analysis. Core genes were extracted from the GFF3 files based on their translated amino acid sequence using a cd-hit cutoff for homologous proteins of 95%, and extracted genes were aligned using Roary (v3.11.2) [79]. The core genome alignment was reduced to only loci harboring polymorphisms using SNP-sites (v2.4.1) [80]. A Maximum Likelihood phylogenetic tree was built using RAxML [77] with the GTR+Gamma model. Statistical branch support was obtained from 100 bootstrap repeats. Phylogenetic trees were visualized from RAxML-generated newick files using TreeGraph 2 (v2.15.0-887 beta) [78]. Branches with bootstrapping support values <70 were collapsed.

Single protein phylogenetic trees were constructed as follows: Nucleotide sequences for genes of interest were extracted from annotated genomes and translated in Geneious (v2019.0.4). Only full length sequences were used to generate single protein phylogenetic trees. Any partial or truncated protein sequences were discarded. Protein sequences were aligned using Muscle (v3.8.425) [74]. Pairwise Muscle alignments were used for tree building with RAxML as described above. Phylogenetic trees were visualized with TreeGraph 2 (v2.15.0-887 beta) [78]. Branches with bootstrapping support values <70 were collapsed.

### T6SS effector and immunity protein typing and subtyping

Effector/immunity protein typing: All sequence manipulations were performed in Geneious (v2019.0.4). All effector and immunity gene sequences were extracted from all identified Aux1 and Accessory Aux1 clusters from all analyzed species. All effectors and immunity nucleotide sequences were translated. Blastp was performed against a custom effector or immunity gene database generated by Kirchberger *et al*., 2017 [24] for each amino acid sequences to identify effector or immunity type (Ex: A, B, C, etc.). For a given sequence, the strongest hit over 30% identity was considered its type [24,26].

Effector/immunity protein subtyping: All effector or immunity amino acid sequences for a given type were aligned with MUSCLE (v3.8.425) [74] to generate a pairwise amino acid identity matrix. Only full-length sequences were used for this analysis. Clusters were determined by inputting protein fasta files to CD-HIT [81,82] with an 80% identity clustering cut off for effector proteins (ex: A1, A2, A3, etc) and a 70% cutoff for immunity genes (ex: A1a, A1b). An 80% identity cutoff for effector proteins is based on experimental data showing a loss of neutralization between two effector/immunity pairs of the same type below 80% identity. Immunity proteins were more variable than effector proteins and thus the identity cutoff was dropped to 70% to be conservative in our typing scheme. Pairwise matrices were used to generate heatmaps using pheatmap (pheatmap, R package pheatmap v1.0.12).

### Alignment of T6SS clusters

Full length T6SS clusters were aligned using the Artemis Comparison Tool (ACT, v18.1.0) [83]. Nucleotide sequences were submitted to NCBI blastn with the “align two or more sequences” option. Alignment files were exported from NCBI and input to ACT along with GenBank format files of the aligned regions.

### Bacterial Strains, Plasmids, and Growth Conditions

*V. cholerae* strains, *V. anguillarum* strains, *E. coli* strains, plasmids, and primers used in this study are listed in Additional File 2: Tables S3 and S4. *E. coli* strain DH5α λpir was used for cloning. All *E. coli* strains were routinely cultured at 37°C in Lysogeny Broth – Lennox (LB) with shaking at 250 rpm. Culture on agar plates was done on LB agar at 28°C (*V. cholerae, V. anguillarum,* and *E. coli).* When required IPTG (for the induction of the T7 polymerase), NaCl (for *V. anguillarum* culture), or antibiotics were added to liquid or agar culture medium at the following concentrations: 100 μM IPTG, 2% w/v NaCl, 100 μg/mL streptomycin, 50 μg/mL rifampicin, 50 μg/mL kanamycin, 100 μg/mL ampicillin, and 30 μg/mL chloramphenicol.

### Bacterial Strain and Plasmid Construction

For all pET vectors, vectors were cut with XhoI and BamHI. Effector or immunity genes were amplified without their stop codon and inserted by Gibson cloning. Primers were modified to ensure that the start codon was in frame with the upstream start codon of the periplasmic localization signal sequence (pelB). For all genes, primers were modified to add a triple glycine linker between the end of the inserted gene and the vector-encoded 6x-His tag.

### Competitive killing assays

Predator *V. cholerae* (streptomycin resistant) and prey *V. anguillarum* (rifampicin resistant) strains were cultured as lawns on lysogeny broth (LB) agar plates with 2% w/v NaCl and selective antibiotics. Plates were incubated for 18-20 hr at 28°C. Predator and prey cells were collected from the overnight lawns with a sterile loop and resuspended in LB + 2% NaCl to an optical density at 600 nm (OD600) of 1. Input cell counts were determined by plating 10-fold serial dilutions of single strain cell suspensions on either LB agar + 2% NaCl + rifampicin or LB agar + 2% NaCl + streptomycin plates. Predator and prey cells were mixed at a 1:1 ratio, and 25 μL of each mixture was spotted on LB agar + 2% NaCl. Competitive mixture spots were incubated for 4 hr at 28°C. Spots were harvested by excision of the underlying agar and vortexing in 1 mL LB + 2% NaCl. For each killing assay, 10-fold serial dilutions were prepared from output cell suspensions, and dilution series were plated on both LB agar + 2% NaCl + rifampicin and LB agar + 2% NaCl + streptomycin plates. Input and output plates were incubated for 18-20 hr at 28°C. Input and output CFU counts were determined for predator and prey strains from each killing assay. Competitive indices were determined as follows: C.I. = [(Output CFU/mL^PREDATOR^/Output CFU/mL^PREY^)/(Input CFU/mL^PREDATOR^/Input CFU/mL^PREY^)].

### Effector/Immunity Co-expression Viability Assays

*E. coli* BL21 (DE3) pLysS strains carrying all pairwise combinations of vectors from the following two lists were generated: (1) pET26b, pET26b-tseL, or pET26b-Aeff^V09^ and (2) pET22b, pET22b-tsiV1, or pET22b-Aimm^V09^. *E. coli* strains were grown (18-20 hr, 37°C, and 250 rpm shaking) in 1 mL LB with selective antibiotics for all three vectors: 30 μg/mL chloramphenicol, 50 μg/mL kanamycin, and 100 μg/mL ampicillin. OD_600_ was calculated, and each overnight culture was normalized to an OD_600_ of 1 in fresh LB. Normalized cell suspensions were spotted (10μL) on LB agar + chloramphenicol + kanamycin + ampicillin both with and without 100 μM IPTG (inducing and non-inducing, respectively). Spot plates were incubated for 18-20 hr at 28°C. Spots from inducing and non-inducing plates were collected by scraping and resuspended in 1 mL LB by vortexing. Output cell counts were determined by plating 10-fold serial dilutions of each cell suspension on LB agar + chloramphenicol + kanamycin + ampicillin. Output dilution plates were incubated for 18-20 hr at 28°C. Viability indices were calculated as follows: Viability Index = [(Output CFU/mL^+IPTG^)/(Output CFU/mL^-IPTG^)].

## DECLARATIONS

### Ethics approval and consent to participate

Not applicable.

### Consent for publication

Not applicable.

### Availability of data and materials

Accession codes for all publicly available genomes analyzed in this study can be found in Additional file 2: Table S1. All analyses are performed with publicly-available tools referenced in the Methods. Custom protein databases referred to in the text are available upon request.

### Authors’ contributions

FJS, PCK, YB, SUP designed the study. FJS and PCK performed analyses. FJS wrote the manuscript. All authors read and approved of the final manuscript.

## Acknowledgements

We thank Hans Rediers and Daniele Provenzano for the strains and reagents necessary for the completion of this study.

## Additional Files

Additional File 1 (.pdf): Supplementary Figures. Figures S1-S7 supporting the data presented in the main manuscript.

Additional File 2 (.pdf): Supplementary Tables. Tables S1-S4 providing information on all genome files, bacterial strains, and primer pairs used to generate the presented data.

Additional File 3 (.xls): T6SS Cluster Information. Table describing the type and location of all T6SS large clusters identified from all analyzed genomes.

Additional File 4 (.xls): *Vang* T6SS Auxiliary Clusters. Table describing the location of all Auxiliary T6SS loci identified from all analyzed *V. anguillarum* genomes.

