## Additional File 1 - Supplemental Figures 1-7 for "Pandemic *Vibrio cholerae* Acquired Competitive Traits from an Environmental *Vibrio* Species"

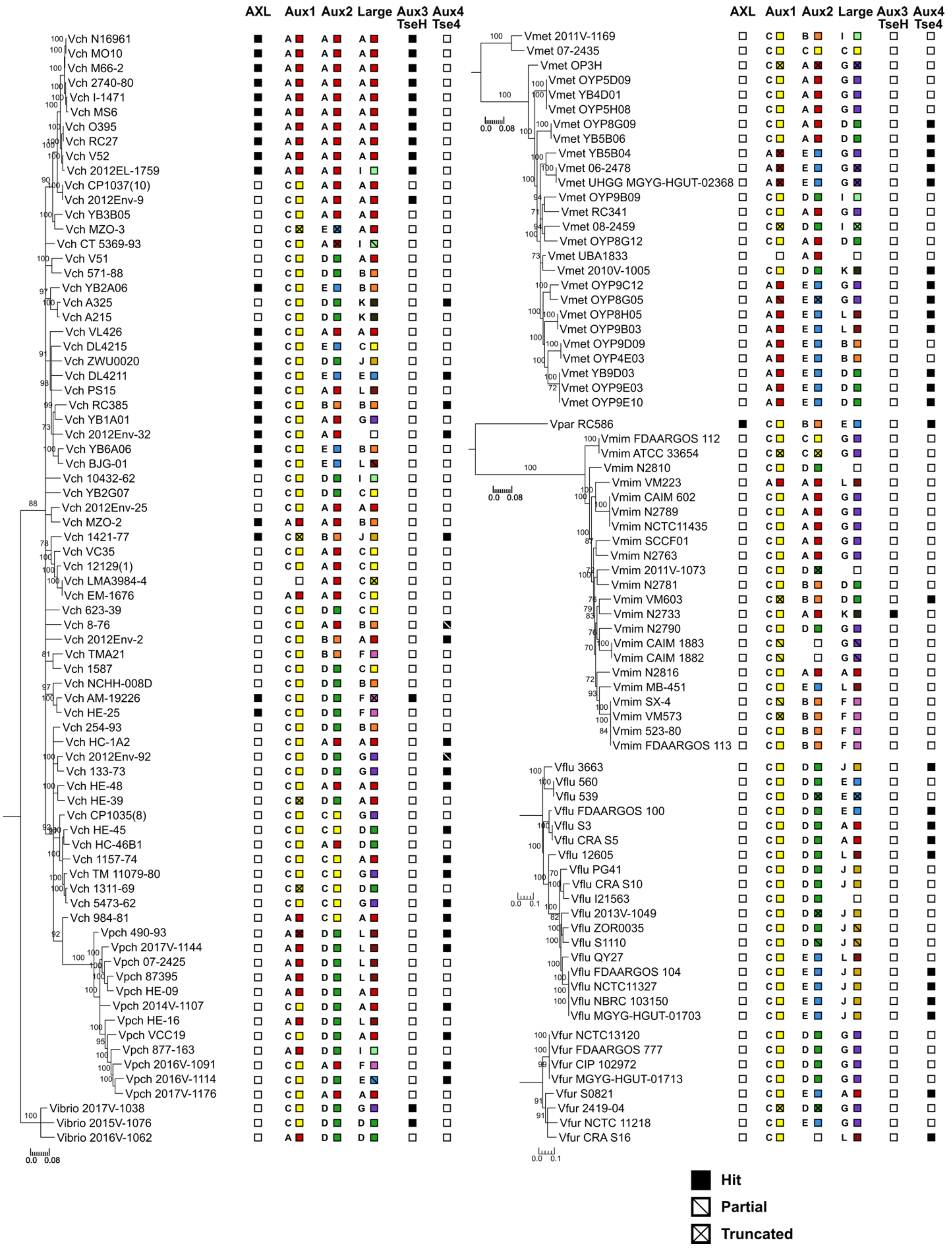

AXL
Aux1
Aux2
Large
Aux3 TseH
Aux4 Tse4
Vch N16961
Vch MO10
Vch M66-2
Vch 2740-80
Vch I-1471
Vch MS6
Vch O395
Vch RC27
Vch V52
Vch 2012EL-1759
Vch CP1037(10)
Vch 2012Env-9
Vch YB3B05
Vch MZO-3
Vch CT 5369-93
Vch V51
Vch 571-88
Vch YB2A06
Vch A325
Vch A215
Vch VL426
Vch DL4215
Vch ZWU0020
Vch DL4211
Vch PS15
Vch RC385
Vch YB1A01
Vch 2012Env-32
Vch YB6A06
Vch BJG-01
Vch 10432-62
Vch YB2G07
Vch 2012Env-25
Vch MZO-2
Vch 1421-77
Vch VC35
Vch 12129(1)
Vch LMA3984-4
Vch EM-1676
Vch 623-39
Vch 8-76
Vch 2012Env-2
Vch TMA21
Vch 1587
Vch NCHH-008D
Vch AM-19226
Vch HE-25
Vch 254-93
Vch HC-1A2
Vch 2012Env-92
Vch 133-73
Vch HE-48
Vch HE-39
Vch CP1035(8)
Vch HE-45
Vch HC-46B1
Vch 1157-74
Vch TM 11079-80
Vch 1311-69
Vch 5473-62
Vch 984-81
Vpch 490-93
Vpch 2017V-1144
Vpch 07-2425
Vpch 87395
Vpch HE-09
Vpch 2014V-1107
Vpch HE-16
Vpch VCC19
Vpch 877-163
Vpch 2016V-1091
Vpch 2016V-1114
Vpch 2017V-1176
Vibrio 2017V-1038
Vibrio 2015V-1076
Vibrio 2016V-1062
0.0 0.08
Vmet 2011V-1169
Vmet 07-2435
Vmet OP3H
Vmet OYP5D09
Vmet YB4D01
Vmet OYP5H08
Vmet OYP8G09
Vmet YB5B06
Vmet YB5B04
Vmet 06-2478
Vmet UHGG MGYG-HGUT-02368
Vmet OYP9B09
Vmet RC341
Vmet 08-2459
Vmet OYP8G12
Vmet UBA1833
Vmet 2010V-1005
Vmet OYP9C12
Vmet OYP8G05
Vmet OYP8H05
Vmet OYP9B03
Vmet OYP9D09
Vmet OYP4E03
Vmet YB9D03
Vmet OYP9E03
Vmet OYP9E10
Vpar RC586
Vmim FDAARGOS 112
Vmim ATCC 33654
Vmim N2810
Vmim VM223
Vmim CAIM 602
Vmim N2789
Vmim NCTC11435
Vmim SCCF01
Vmim N2763
Vmim 2011V-1073
Vmim N2781
Vmim VM603
Vmim N2733
Vmim N2790
Vmim CAIM 1883
Vmim CAIM 1882
Vmim N2816
Vmim MB-451
Vmim SX-4
Vmim VM573
Vmim 523-80
Vmim FDAARGOS 113
Vflu 3663
Vflu 560
Vflu 539
Vflu FDAARGOS 100
Vflu S3
Vflu CRA S5
Vflu 12605
Vflu PG41
Vflu CRA S10
Vflu I21563
Vflu 2013V-1049
Vflu ZOR0035
Vflu S1110
Vflu QY27
Vflu FDAARGOS 104
Vflu NCTC11327
Vflu NBRC 103150
Vflu MGYG-HGUT-01703
Vfur NCTC13120
Vfur FDAARGOS 777
Vfur CIP 102972
Vfur MGYG-HGUT-01713
Vfur S0821
Vfur 2419-04
Vfur NCTC 11218
Vfur CRA S16
0.0 0.1
Hit
Partial
Truncated

**Figure S1. T6SS cluster and effector/immunity type distribution across *Vch, Vpch, Vmet, Vpar*, *Vmim, Vflu, and Vfur strains.***
Species trees for *Vch*, *Vpch*, *Vmet*, *Vpar, Vmim*, *Vflu*, and *Vfur*. One maximum-likelihood tree based on core-genome SNP sites for each species or group of species. Bootstrapping support values are shown next to corresponding nodes. All nodes with bootstrapping support values below 70 are collapsed. Columns next to trees represent T6SS effectors at different clusters from the *Vch* population. Filled boxes represent presence. Empty boxes represent absence. Partial or truncated hits are indicated by slashes. For effectors with multiple types, both letter and corresponding color (Fig. 1c) of the present effector type are shown. All scale bars represent Substitutions/SNP site.

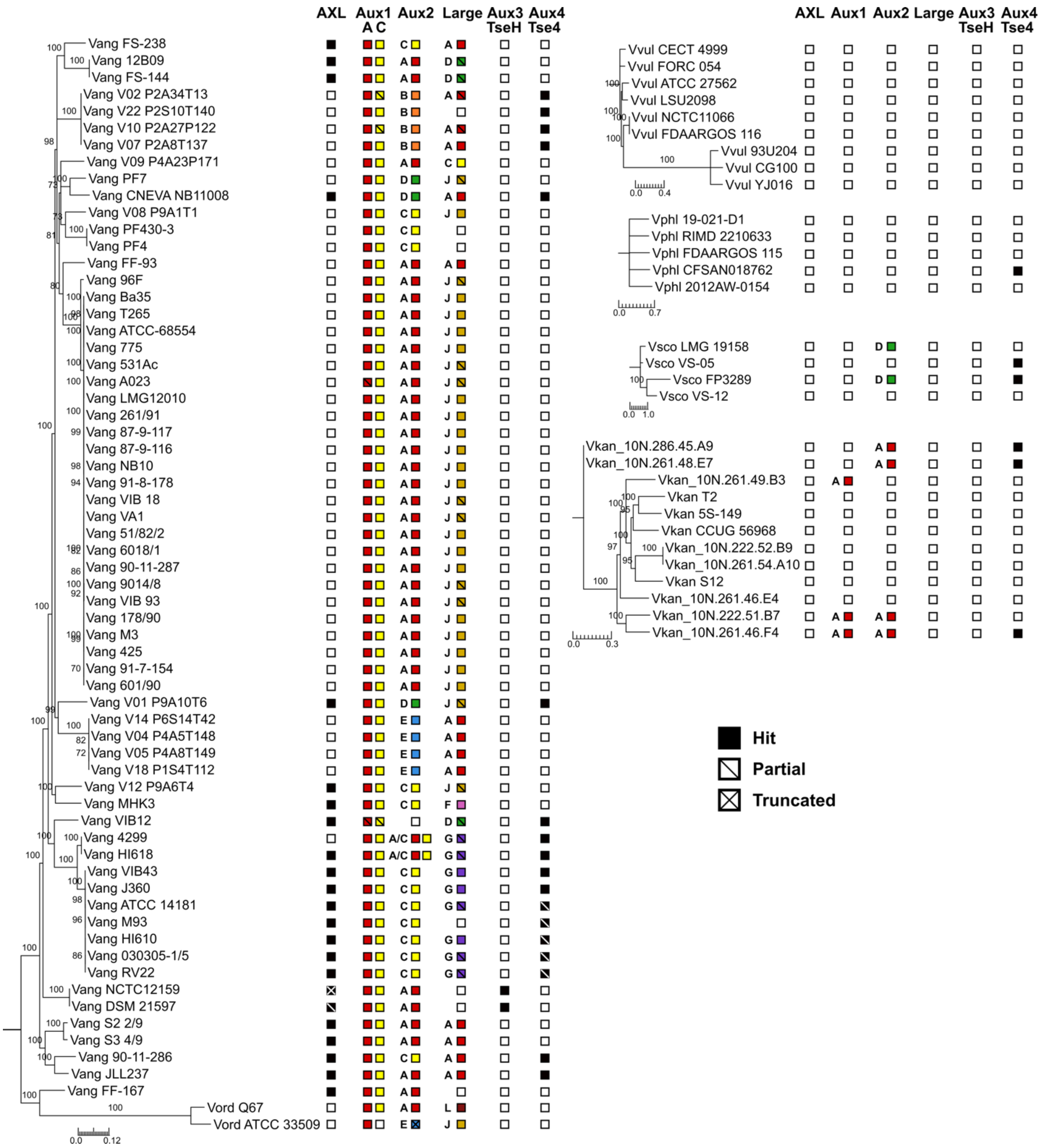

**Figure S2. T6SS cluster and effector/immunity type distribution across *Vang, Vord, Vvul, Vphl, Vsco, and Vkan strains.***

Species trees for *Vang*, *Vord*, *Vvul*, *Vphl*, *Vsco*, and *Vkan*. One maximum-likelihood tree based on core-genome SNP sites for each species. Bootstrapping support values are shown next to corresponding nodes. All nodes with bootstrapping support values below 70 are collapsed. Columns next to trees represent T6SS effectors at different clusters from the *Vch* population. Filled boxes represent presence. Empty boxes represent absence. Partial or truncated hits are indicated by slashes. For effectors with multiple types, both letter and corresponding color (Fig. 1c) of the present effector type are shown. All scale bars represent Substitutions/SNP site.

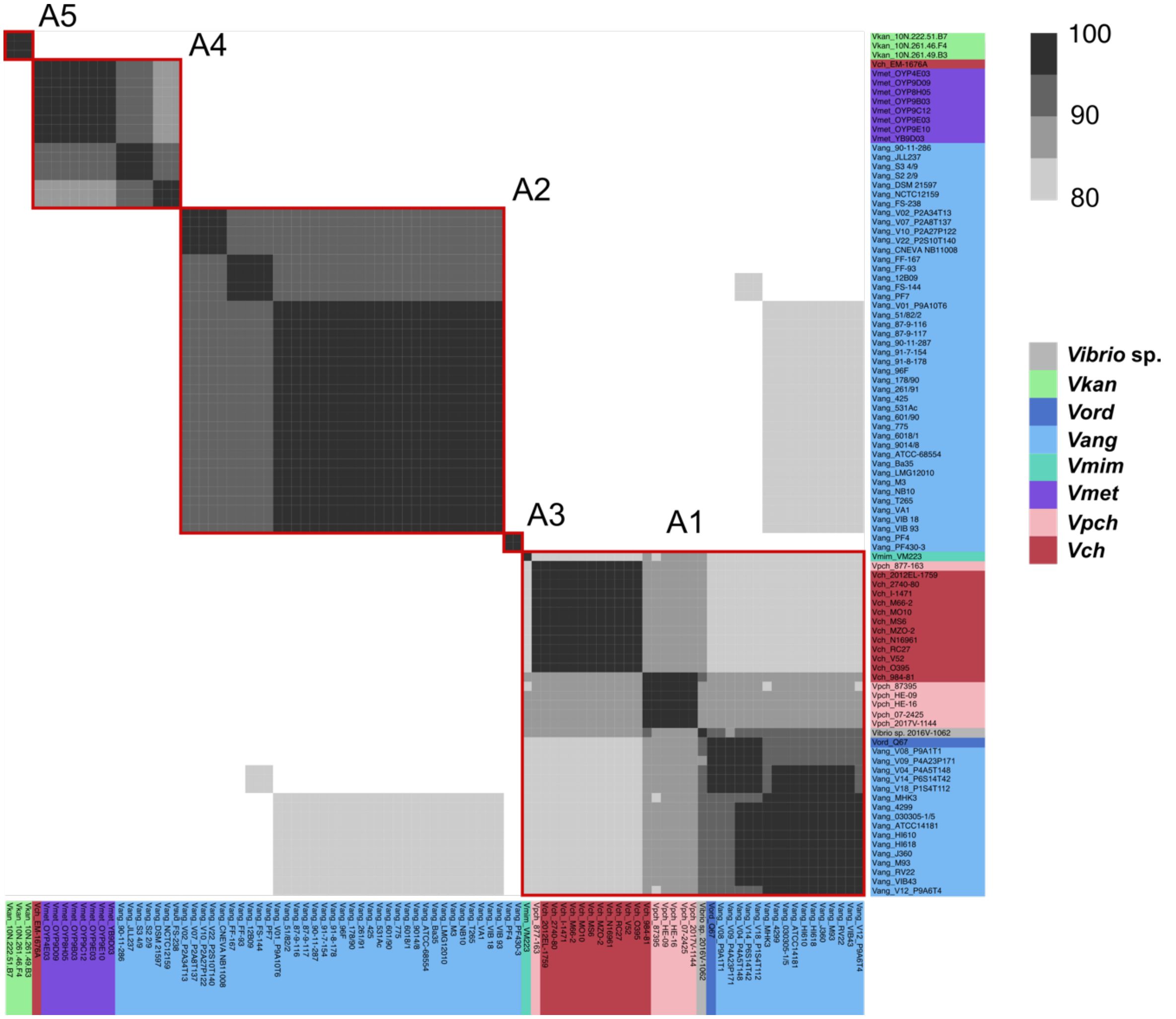


**Figure S3. Aux1 A-type effector proteins are divided into subtypes.**
Pairwise amino acid identity heatmap of Aux1 A effector proteins (TseL) from *Vch*, *Vpch*, *Vmet*, *Vmim*, *Vang*, *Vord*, *Vkan*, and Vibrio sp. closely related to *Vch*. Heatmap is restricted to 80%-100% identity. Distinct clusters called by CD-HIT with a 80% identity cutoff are indicated by red boxes.

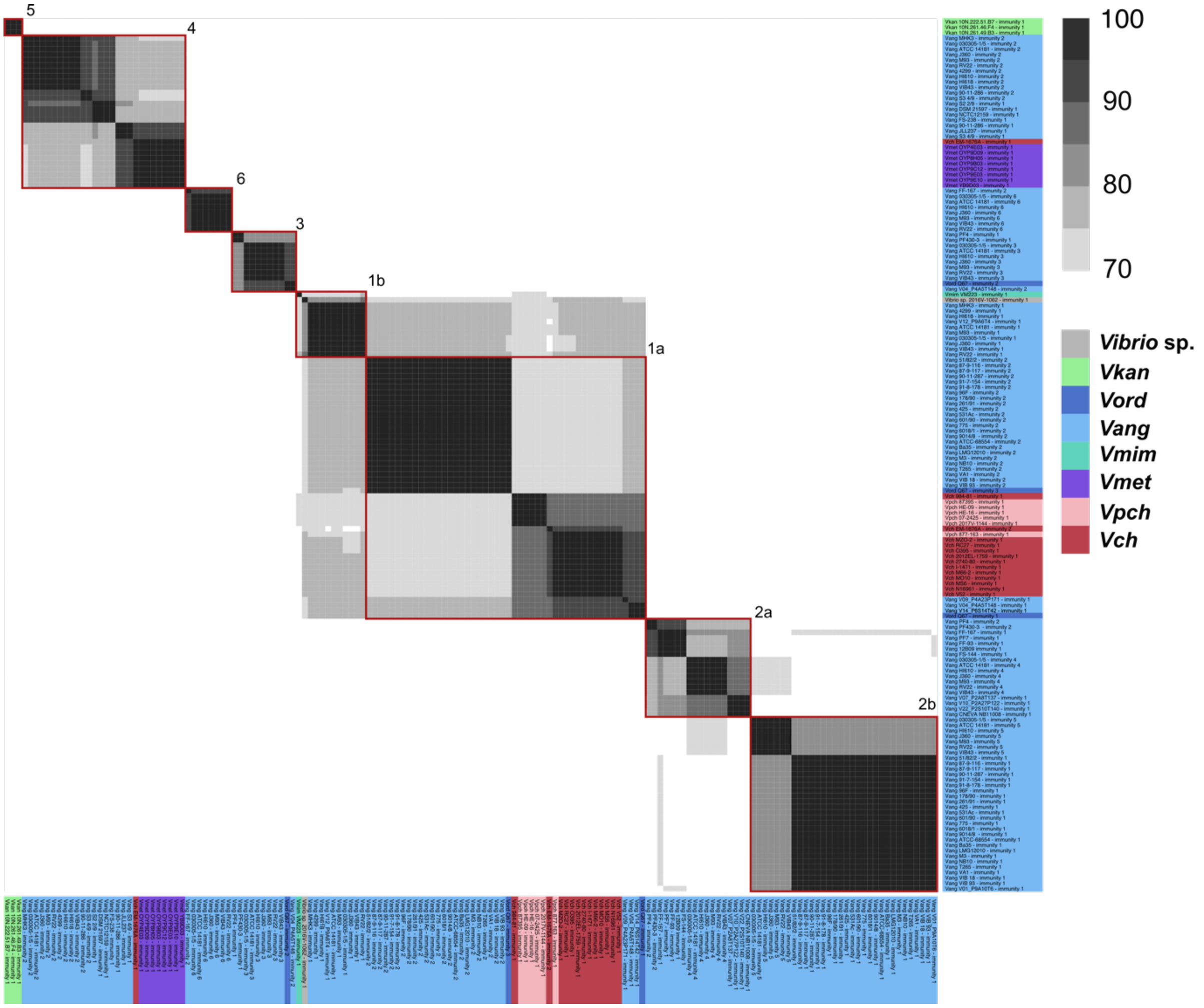


**Figure S4. Aux1 A-type immunity proteins are further divided beyond subtype.** Pairwise amino acid identity heatmap of Aux1 A immunity proteins (TsiV1) from *Vch*, *Vpch*, *Vmet*, *Vmim*, *Vang*, *Vord*, *V. kan*, and Vibrio sp. closely related to *Vch*. Heatmap is restricted to 70%-100% identity. Distinct clusters called by CD-HIT with a 70% identity cutoff are indicated by red boxes.

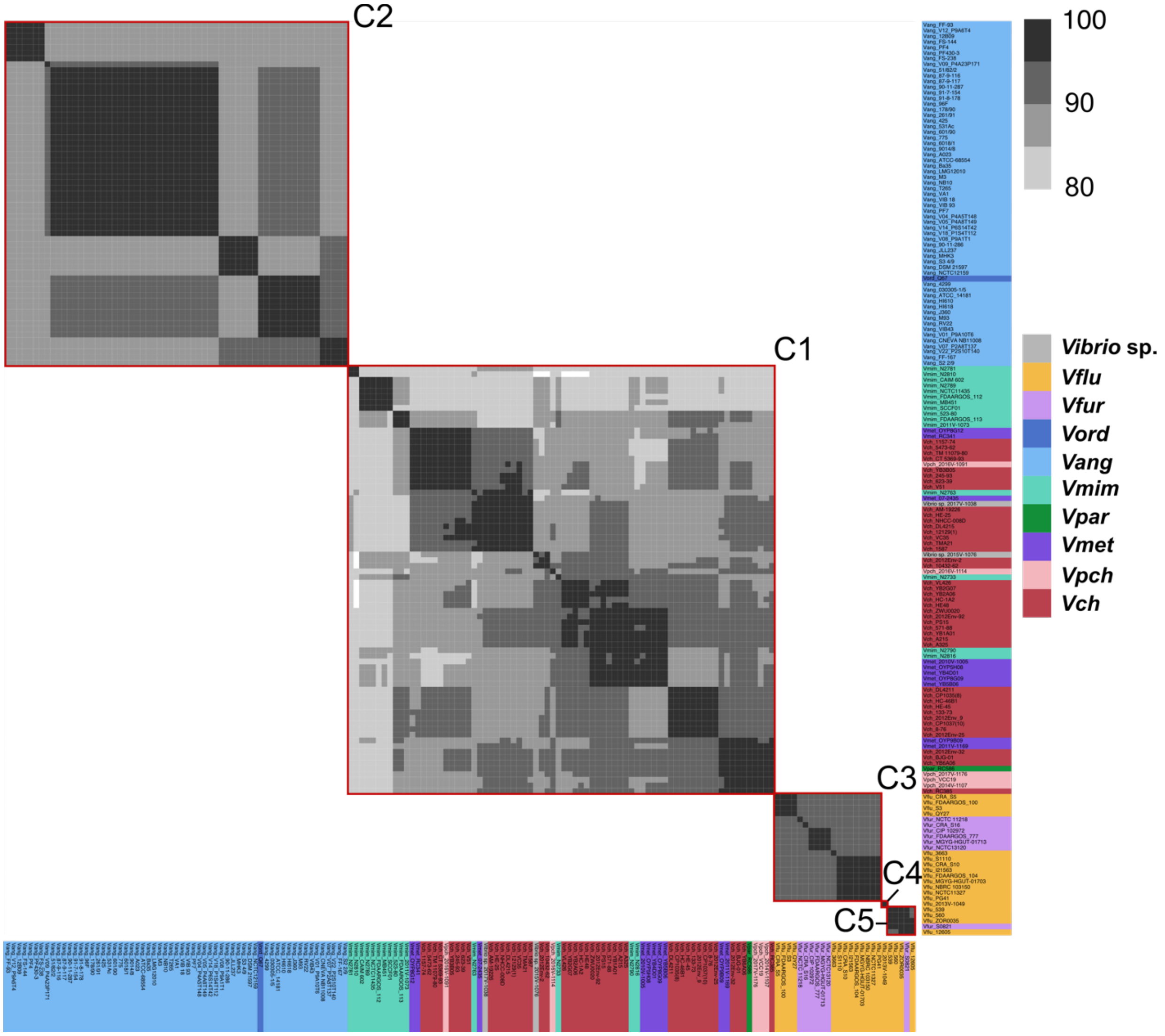


**Figure S5. Aux1 C-type effector proteins are divided into subtypes.**
Pairwise amino acid identity heatmap of Aux1 C effector proteins from *Vch*, *Vpch*, *Vmet*, *Vpar*, *Vmim*, *Vang*, *Vord*, *Vfur*, *Vflu* , and Vibrio sp. closely related to *Vch*. Heatmap is restricted to 80%-100% identity. Distinct clusters called by CD-HIT with a 80% identity cutoff are indicated by red boxes.

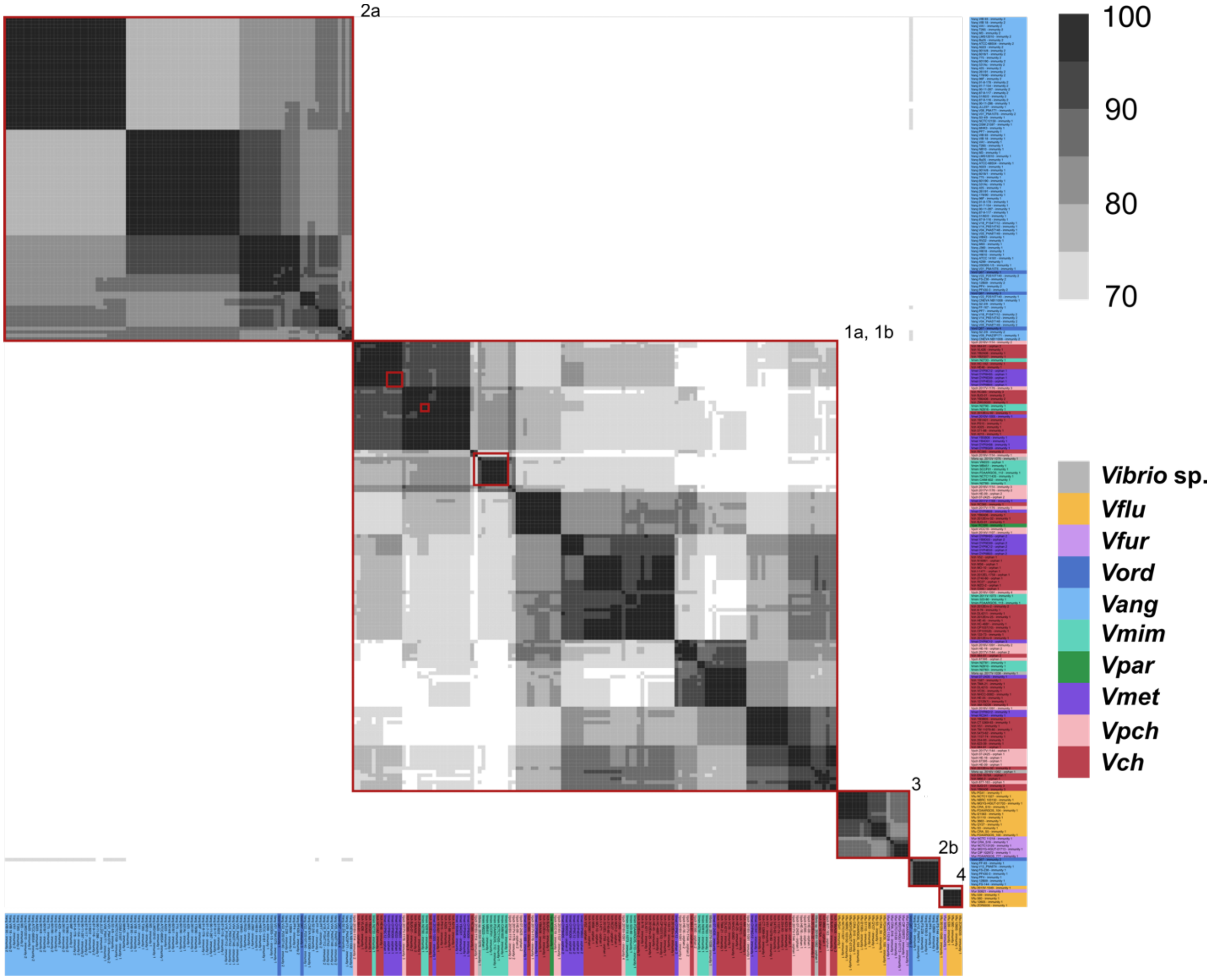


**Figure S6. Aux1 C-type immunity proteins are further divided beyond subtype.**
Pairwise amino acid identity heatmap of Aux1 C immunity proteins from *Vch*, *Vpch*, *Vmet*, *Vpar*, *Vmim*, *Vang*, *Vord*, *Vfur*, *Vflu* , and Vibrio sp. closely related to *Vch*. Heatmap is restricted to 70%-100% identity. Distinct clusters called by CD-HIT with a 70% identity cutoff are indicated by red boxes.

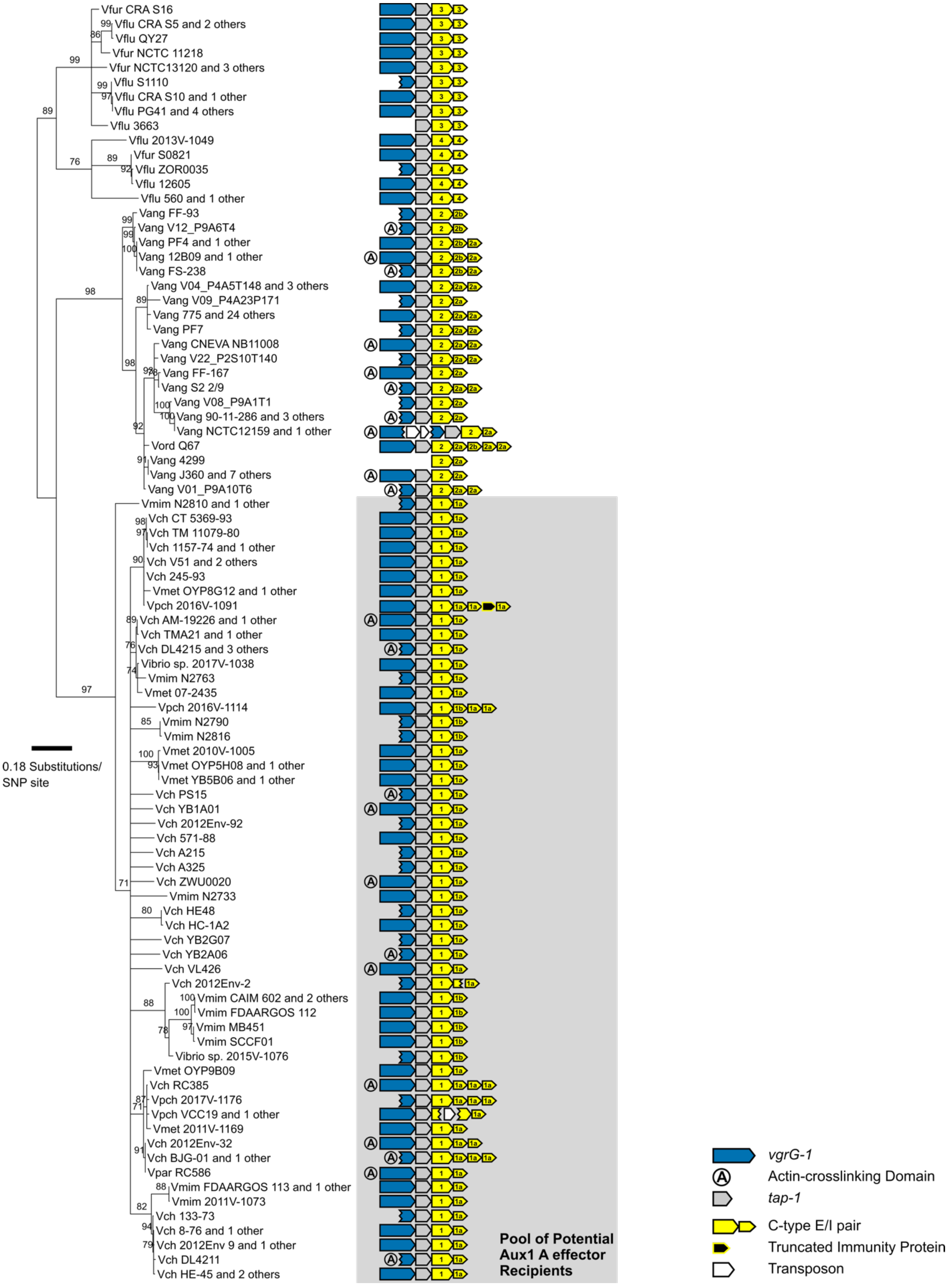
Vfur CRA S16
Vflu CRA S5 and 2 others
Vflu QY27
Vfur NCTC 11218
Vfur NCTC13120 and 3 others
Vflu S1110
Vflu CRA S10 and 1 other
Vflu PG41 and 4 others
Vflu 3663
Vflu 2013V-1049
Vfur S0821
Vflu ZOR0035
Vflu 12605
Vflu 560 and 1 other
Vang FF-93
Vang V12_P9A6T4
Vang PF4 and 1 other
Vang 12B09 and 1 other
Vang FS-238
Vang V04_P4A5T148 and 3 others
Vang V09_P4A23P171
Vang 775 and 24 others
Vang PF7
Vang CNEVA NB11008
Vang V22_P2S10T140
Vang FF-167
Vang S2 2/9
Vang V08_P9A1T1
Vang 90-11-286 and 3 others
Vang NCTC12159 and 1 other
Vord Q67
Vang 4299
Vang J360 and 7 others
Vang V01_P9A10T6
Vmim N2810 and 1 other
Vch CT 5369-93
Vch TM 11079-80
Vch 1157-74 and 1 other
Vch V51 and 2 others
Vch 245-93
Vmet OYP8G12 and 1 other
Vpch 2016V-1091
Vch AM-19226 and 1 other
Vch TMA21 and 1 other
Vch DL4215 and 3 others
Vibrio sp. 2017V-1038
Vmim N2763
Vmet 07-2435
Vpch 2016V-1114
Vmim N2790
Vmim N2816
Vmet 2010V-1005
Vmet OYP5H08 and 1 other
Vmet YB5B06 and 1 other
Vch PS15
Vch YB1A01
Vch 2012Env-92
Vch 571-88
Vch A215
Vch A325
Vch ZWU0020
Vmim N2733
Vch HE48
Vch HC-1A2
Vch YB2G07
Vch YB2A06
Vch VL426
Vch 2012Env-2
Vmim CAIM 602 and 2 others
Vmim FDAARGOS 112
Vmim MB451
Vmim SCCF01
Vibrio sp. 2015V-1076
Vmet OYP9B09
Vch RC385
Vpch 2017V-1176
Vpch VCC19 and 1 other
Vmet 2011V-1169
Vch 2012Env-32
Vch BJG-01 and 1 other
Vpar RC586
Vmim FDAARGOS 113 and 1 other
Vmim 2011V-1073
Vch 133-73
Vch 8-76 and 1 other
Vch 2012Env 9 and 1 other
Vch DL4211
Vch HE-45 and 2 others
0.18 Substitutions/
SNP site
Pool of Potential
Aux1 A effector
Recipients
vgrG-1
Actin-crosslinking Domain
tap-1
C-type E/I pair
Truncated Immunity Protein
Transposon

**Figure S7. Aux1 C effector/Immunity subtypes are not horizontally transferred between clades.**
Maximum-likelihood tree of identified Aux1 C-type effectors from strains lacking an AccAux1 cluster or an A-type effector at Aux1. Bootstrapping support values for each node are shown. Aux1 schematic for each strain is shown on the right. Each effector/immunity cassette is labeled with its associated subtype. Grey box indicates group of potential recipient strains involved in transfer of the A-type effector from *Vang* and the generation of an orphan immunity gene in *Vch*.
