## Additional File 2 - Supplemental Tables 1-3 for "Pandemic *Vibrio cholerae* Acquired Competitive Traits from an Environmental *Vibrio* Species"

**Table S1. Vibrio genomes analyzed in this study.**

| Sequence Name | RefSeq or GenBank Accession |
|---|---|
| Vibrio anguillarum strain 12B09 | GCF_000287135.2 |
| Vibrio anguillarum strain 51/82/2 | GCA_001989855.1 |
| Vibrio anguillarum strain 87-9-116 | GCF_002211505.1 |
| Vibrio anguillarum strain 87-9-117 | GCA_001989715.1 |
| Vibrio anguillarum strain 90-11-286 | GCF_001660505.1 |
| Vibrio anguillarum strain 90-11-287 | GCA_001990025.1 |
| Vibrio anguillarum strain 91-7-154 | GCA_001990045.1 |
| Vibrio anguillarum strain 91-8-178 | GCA_001989735.1 |
| Vibrio anguillarum strain 96F | GCF_000257165.1 |
| Vibrio anguillarum strain 178/90 | GCA_001989755.1 |
| Vibrio anguillarum strain 261/91 | GCA_001989775.1 |
| Vibrio anguillarum strain 425 | GCF_003031205.1 |
| Vibrio anguillarum strain 531Ac | GCF_008107435.1 |
| Vibrio anguillarum strain 601/90 | GCA_001990065.1 |
| Vibrio anguillarum strain 775 | GCF_000217675.1 |
| Vibrio anguillarum strain 4299 | GCA_001989655.1 |
| Vibrio anguillarum strain 6018/1 | GCA_001990085.1 |
| Vibrio anguillarum strain 9014/8 | GCA_001989915.1 |
| Vibrio anguillarum strain 030305-1/5 | GCF_003709525.1 |
| Vibrio anguillarum strain A023 | GCA_001989675.1 |
| Vibrio anguillarum strain ATCC 14181 | GCF_001718015.1 |
| Vibrio anguillarum strain ATCC-68554 | GCF_002291265.1 |
| Vibrio anguillarum strain Ba35 | GCA_001989795.1 |
| Vibrio anguillarum strain CNEVA NB11008 | GCF_002212025.1 |
| Vibrio anguillarum strain DSM 21597 | GCA_001989995.1 |
| Vibrio anguillarum strain FF-93 | GCF_000287095.2 |
| Vibrio anguillarum strain FF-167 | GCF_000287075.2 |
| Vibrio anguillarum strain FS-144 | GCF_000287115.2 |
| Vibrio anguillarum strain FS-238 | GCF_000287155.2 |
| Vibrio anguillarum strain HI610 | GCA_001989835.1 |
| Vibrio anguillarum strain HI618 | GCF_002078035.1 |
| Vibrio anguillarum strain J360 | GCF_003399575.2 |
| Vibrio anguillarum strain JLL237 | GCF_002211985.1 |
| Vibrio anguillarum strain LMG12010 | GCA_001989875.1 |
| Vibrio anguillarum strain M3 | GCF_000462975.1 |
| Vibrio anguillarum strain M93 | GCF_002901125.1 |

| | |
|---|---|
| Vibrio anguillarum strain MHK3 | GCF_003595585.1 |
| Vibrio anguillarum strain NB10 | GCF_000786425.1 |
| Vibrio anguillarum strain NCTC12159 | GCF_900452855.1 |
| Vibrio anguillarum strain PF4 | GCA_002813835.1 |
| Vibrio anguillarum strain PF7 | GCA_001997225.1 |
| Vibrio anguillarum strain PF430-3 | GCA_001989695.1 |
| Vibrio anguillarum strain RV22 | GCF_000257185.1 |
| Vibrio anguillarum strain S2 2/9 | GCA_001989895.1 |
| Vibrio anguillarum strain S3 4/9 | GCF_002212005.1 |
| Vibrio anguillarum strain T265 | GCA_001989815.1 |
| Vibrio anguillarum strain V01_P9A10T6 | GCF_002995435.1 |
| Vibrio anguillarum strain V02_P2A34T13 | GCF_003006755.1 |
| Vibrio anguillarum strain V04_P4A5T148 | GCF_002241075.1 |
| Vibrio anguillarum strain V05_P4A8T149 | GCF_002241155.1 |
| Vibrio anguillarum strain V07_P2A8T137 | GCF_002241165.1 |
| Vibrio anguillarum strain V08_P9A1T1 | GCF_002241195.1 |
| Vibrio anguillarum strain V09_P4A23P171 | GCF_002241235.1 |
| Vibrio anguillarum strain V10_P2A27P122 | GCF_002241255.1 |
| Vibrio anguillarum strain V12_P9A6T4 | GCF_002241275.1 |
| Vibrio anguillarum strain V14_P6S14T42 | GCF_002241285.1 |
| Vibrio anguillarum strain V18_P1S4T112 | GCF_002241365.1 |
| Vibrio anguillarum strain V22_P2S10T140 | GCF_009905095.1 |
| Vibrio anguillarum strain VA1 | GCA_001990105.1 |
| Vibrio anguillarum strain VIB12 | GCF_002310335.1 |
| Vibrio anguillarum strain VIB43 | GCF_002287545.1 |
| Vibrio anguillarum strain VIB 18 | GCA_001998845.1 |
| Vibrio anguillarum strain VIB 93 | GCA_001990125.1 |
| Vibrio cholerae strain 8-76 | GCF_000736935.1 |
| Vibrio cholerae strain 133-73 | GCF_000736765.1 |
| Vibrio cholerae strain 254-93 | GCF_000737025.1 |
| Vibrio cholerae strain 571-88 | GCF_000736945.1 |
| Vibrio cholerae strain 623-39 | GCF_000154005.2 |
| Vibrio cholerae strain 984-81 | GCF_000736775.1 |
| Vibrio cholerae strain 1157-74 | GCF_000736875.1 |
| Vibrio cholerae strain 1311-69 | GCF_000736855.1 |
| Vibrio cholerae strain 1421-77 | GCF_000736785.1 |
| Vibrio cholerae strain 1587 | GCF_000168895.2 |
| Vibrio cholerae strain 2012EL-1759 | GCF_000710155.1 |
| Vibrio cholerae strain 2012Env-2 | GCF_000788495.1 |

| | |
|---|---|
| Vibrio cholerae strain 2012Env-9 | GCF_000788715.2 |
| Vibrio cholerae strain 2012Env-25 | GCF_000788515.1 |
| Vibrio cholerae strain 2012Env-32 | GCF_000788675.1 |
| Vibrio cholerae strain 2012Env-92 | GCF_000788755.1 |
| Vibrio cholerae strain 2740-80 | GCF_000168915.2 |
| Vibrio cholerae strain 5473-62 | GCF_000736795.1 |
| Vibrio cholerae strain 10432-62 | GCF_000969265.1 |
| Vibrio cholerae strain 12129(1) | GCF_000174115.1 |
| Vibrio cholerae strain A215 | GCF_001259995.1 |
| Vibrio cholerae strain A325 | GCF_001254095.1 |
| Vibrio cholerae strain AM-19226 | GCF_000153785.2 |
| Vibrio cholerae strain BJG-01 | GCF_000221465.1 |
| Vibrio cholerae strain CP1035(8) | GCF_000304915.2 |
| Vibrio cholerae strain CP1037(10) | GCF_000302965.1 |
| Vibrio cholerae strain CT 5369-93 | GCF_000176455.1 |
| Vibrio cholerae strain DL4211 | GCF_001953365.1 |
| Vibrio cholerae strain DL4215 | GCF_001953375.1 |
| Vibrio cholerae strain EM-1676A | GCF_000348345.2 |
| Vibrio cholerae strain HC-1A2 | GCF_000304775.1 |
| Vibrio cholerae strain HC-46B1 | GCF_000305605.1 |
| Vibrio cholerae strain HE-25 | GCF_000279265.1 |
| Vibrio cholerae strain HE-39 | GCF_000220765.2 |
| Vibrio cholerae strain HE-45 | GCF_000279285.1 |
| Vibrio cholerae strain HE-48 | GCF_000220785.1 |
| Vibrio cholerae strain I-1471 | GCF_000818865.1 |
| Vibrio cholerae strain LMA3984-4 | GCF_000195065.1 |
| Vibrio cholerae strain MO10 | GCF_000152425.1 |
| Vibrio cholerae strain M66-2 | GCF_000021605.1 |
| Vibrio cholerae strain MS6 | GCF_000829215.1 |
| Vibrio cholerae strain MZO-2 | GCF_000153985.2 |
| Vibrio cholerae strain MZO-3 | GCF_000168935.2 |
| Vibrio cholerae strain N16961 | GCF_000006745.1 |
| Vibrio cholerae strain NHCC-008D | GCF_000348425.2 |
| Vibrio cholerae strain O395 | GCF_000021625.1 |
| Vibrio cholerae strain PS15 | GCF_000318075.1 |
| Vibrio cholerae strain RC27 | GCF_000176395.1 |
| Vibrio cholerae strain RC385 | GCF_000152445.1 |
| Vibrio cholerae strain TM 11079-80 | GCF_000174255.1 |
| Vibrio cholerae strain TMA21 | GCF_000174295.1 |

| | |
|---|---|
| Vibrio cholerae strain V51 | GCF_000152465.2 |
| Vibrio cholerae strain V52 | GCF_000167935.2 |
| Vibrio cholerae strain VC35 | GCF_000299495.2 |
| Vibrio cholerae strain VL426 | GCF_000174235.1 |
| Vibrio cholerae strain YB1A01 | GCF_001402185.1 |
| Vibrio cholerae strain YB2A06 | GCF_001402375.1 |
| Vibrio cholerae strain YB2G07 | GCF_001402425.1 |
| Vibrio cholerae strain YB3B05 | GCF_001402545.1 |
| Vibrio cholerae strain YB6A06 | GCF_001402445.1 |
| Vibrio cholerae strain ZWU0020 | GCF_000812045.1 |
| Vibrio fluvialis strain 539 | GCF_000760625.1 |
| Vibrio fluvialis strain 560 | GCF_000754645.1 |
| Vibrio fluvialis strain 2013V-1049 | GCF_009665355.1 |
| Vibrio fluvialis strain 3663 | GCF_000931495.1 |
| Vibrio fluvialis strain 12605 | GCF_001952955.1 |
| Vibrio fluvialis strain CRA_S5 | GCF_007050325.1 |
| Vibrio fluvialis strain CRA_S10 | GCF_007050345.1 |
| Vibrio fluvialis strain FDAARGOS_100 | GCF_002953375.1 |
| Vibrio fluvialis strain FDAARGOS_104 | GCF_001558415.2 |
| Vibrio fluvialis strain I21563 | GCF_000418995.1 |
| Vibrio fluvialis strain MGYG-HGUT-01703 | GCF_902377575.1 |
| Vibrio fluvialis strain NBRC 103150 | GCF_001598835.1 |
| Vibrio fluvialis strain NCTC11327 | GCF_900460245.1 |
| Vibrio fluvialis strain PG41 | GCF_000417625.1 |
| Vibrio fluvialis strain QY27 | GCF_002796765.1 |
| Vibrio fluvialis strain S3 | GCF_006381955.1 |
| Vibrio fluvialis strain S1110 | GCF_001418705.1 |
| Vibrio fluvialis strain ZOR0035 | GCF_000799015.1 |
| Vibrio furnissii strain 2419-04 | GCF_009665335.1 |
| Vibrio furnissii strain CIP 102972 | GCF_000176175.1 |
| Vibrio furnissii strain CRA_S16 | GCF_007050385.1 |
| Vibrio furnissii strain FDAARGOS_777 | GCF_006364355.1 |
| Vibrio furnissii strain MGYG-HGUT-01713 | GCF_902377635.1 |
| Vibrio furnissii strain NCTC13120 | GCF_900460225.1 |
| Vibrio furnissii strain NCTC 11218 | GCF_000184325.1 |
| Vibrio furnissii strain S0821 | GCF_001418695.1 |
| Vibrio sp. 2015V-1076 | GCF_003311815.1 |
| Vibrio sp. 2016V-1062 | GCF_003311825.1 |
| Vibrio sp. 2017V-1038 | GCF_003311805.1 |

| | |
|---|---|
| Vibrio parilis RC586 | GCF_000176715.1 |
| Vibrio kanaloae strain 5S-149 | GCF_000272165.2 |
| Vibrio kanaloae strain 10N.222.51.B7 | GCF_005146725.1 |
| Vibrio kanaloae strain 10N.222.52.B9 | GCF_005146545.1 |
| Vibrio kanaloae strain 10N.261.46.E4 | GCF_005146415.1 |
| Vibrio kanaloae strain 10N.261.46.F4 | GCF_005146515.1 |
| Vibrio kanaloae strain 10N.261.48.E7 | GCF_005146445.1 |
| Vibrio kanaloae strain 10N.261.49.B3 | GCF_002876865.1 |
| Vibrio kanaloae strain 10N.261.54.A10 | GCF_005145845.1 |
| Vibrio kanaloae strain 10N.286.45.A9 | GCF_005145785.1 |
| Vibrio kanaloae strain CCUG 56968 | GCF_008801285.1 |
| Vibrio kanaloae strain S12 | GCF_007858925.1 |
| Vibrio kanaloae strain T2 | GCF_007858815.1 |
| Vibrio metoecus strain 06-2478 | GCF_001402155.1 |
| Vibrio metoecus strain 07-2435 | GCF_001402165.1 |
| Vibrio metoecus strain 08-2459 | GCF_009665275.1 |
| Vibrio metoecus strain 2010V-1005 | GCF_001402685.1 |
| Vibrio metoecus strain 2011V-1169 | GCF_009665255.1 |
| Vibrio metoecus strain OP3H | GCF_000696385.1 |
| Vibrio metoecus strain OYP4E03 | GCF_002284005.1 |
| Vibrio metoecus strain OYP5D09 | GCF_002283995.1 |
| Vibrio metoecus strain OYP5H08 | GCF_002283965.1 |
| Vibrio metoecus strain OYP8G05 | GCF_002283955.1 |
| Vibrio metoecus strain OYP8G09 | GCF_002284045.1 |
| Vibrio metoecus strain OYP8G12 | GCF_002284025.1 |
| Vibrio metoecus strain OYP8H05 | GCF_002283915.1 |
| Vibrio metoecus strain OYP9B03 | GCF_002283905.1 |
| Vibrio metoecus strain OYP9B09 | GCF_002283845.1 |
| Vibrio metoecus strain OYP9C12 | GCF_002283835.1 |
| Vibrio metoecus strain OYP9D09 | GCF_002283805.1 |
| Vibrio metoecus strain OYP9E03 | GCF_002283895.1 |
| Vibrio metoecus strain OYP9E10 | GCF_002283855.1 |
| Vibrio metoecus strain RC341 | GCF_000176215.1 |
| Vibrio metoecus strain UBA1833 | GCA_002339045.1 |
| Vibrio metoecus strain UHGG_MGYG-HGUT-02368 | GCF_902386355.1 |
| Vibrio metoecus strain YB4D01 | GCF_001402495.1 |
| Vibrio metoecus strain YB5B04 | GCF_001402675.1 |
| Vibrio metoecus strain YB5B06 | GCF_001402345.1 |

| | |
|---|---|
| Vibrio metoecus strain YB9D03 | GCF_001402515.1 |
| Vibrio mimicus strain 523-80 | GCF_000736955.1 |
| Vibrio mimicus strain 2011V-1073 | GCF_009665195.1 |
| Vibrio mimicus strain ATCC 33654 | GCF_008464965.1 |
| Vibrio mimicus strain CAIM 602 | GCF_000338875.1 |
| Vibrio mimicus strain CAIM 1882 | GCF_000473785.1 |
| Vibrio mimicus strain CAIM 1883 | GCF_000473825.1 |
| Vibrio mimicus strain FDAARGOS_112 | GCF_001558475.2 |
| Vibrio mimicus strain FDAARGOS_113 | GCF_001471395.2 |
| Vibrio mimicus strain MB-451 | GCF_000176375.1 |
| Vibrio mimicus strain N2733 | GCF_008084745.1 |
| Vibrio mimicus strain N2763 | GCF_008084405.1 |
| Vibrio mimicus strain N2781 | GCF_008083965.1 |
| Vibrio mimicus strain N2789 | GCF_008083745.1 |
| Vibrio mimicus strain N2790 | GCF_008083775.1 |
| Vibrio mimicus strain N2810 | GCF_008083535.1 |
| Vibrio mimicus strain N2816 | GCF_008083465.1 |
| Vibrio mimicus strain NCTC11435 | GCF_900460385.1 |
| Vibrio mimicus strain SCCF01 | GCF_001767355.1 |
| Vibrio mimicus strain SX-4 | GCF_000222145.1 |
| Vibrio mimicus strain VM223 | GCF_000176415.1 |
| Vibrio mimicus strain VM573 | GCF_000175995.1 |
| Vibrio mimicus strain VM603 | GCF_000175975.1 |
| Vibrio ordalii strain ATCC 33509 | GCF_000257205.1 |
| Vibrio ordalii strain Q67 | GCA_002257545.1 |
| Vibrio paracholerae strain 07-2425 | GCF_003311905.1 |
| Vibrio paracholerae strain 2014V-1107 | GCF_003311945.1 |
| Vibrio paracholerae strain 2016V-1091 | GCF_003312065.1 |
| Vibrio paracholerae strain 2016V-1114 | GCF_003312085.1 |
| Vibrio paracholerae strain 2017V-1176 | GCF_003312095.1 |
| Vibrio paracholerae strain 2017V-1144 | GCF_003312015.1 |
| Vibrio paracholerae strain 490-93 | GCF_000737015.1 |
| Vibrio paracholerae strain 877-163 | GCF_001402745.1 |
| Vibrio paracholerae strain 87395 | GCF_000348085.2 |
| Vibrio paracholerae strain HE-09 | GCF_000221405.1 |
| Vibrio paracholerae strain HE-16 | GCF_000303085.1 |
| Vibrio paracholerae strain VCC19 | GCF_000438805.2 |
| Vibrio parahaemolyticus strain 19-021-D1 | GCF_009734325.1 |

| | |
|---|---|
| Vibrio parahaemolyticus strain 2012AW-0154 | GCF_009665495.1 |
| Vibrio parahaemolyticus strain CFSAN018762 | GCF_001696155.1 |
| Vibrio parahaemolyticus strain FDAARGOS_115 | GCF_001558495.2 |
| Vibrio parahaemolyticus strain RIMD 2210633 | GCF_000196095.1 |
| Vibrio scophthalmi strain FP3289 | GCF_001723385.1 |
| Vibrio scophthalmi strain LMG 19158 | GCF_000222585.1 |
| Vibrio scophthalmi strain VS-05 | GCF_001687805.1 |
| Vibrio scophthalmi strain VS-12 | GCF_001685465.1 |
| Vibrio vulnificus strain 93U204 | GCF_000746665.1 |
| Vibrio vulnificus strain ATCC 27562 | GCF_002224265.1 |
| Vibrio vulnificus strain CECT 4999 | GCF_002215135.1 |
| Vibrio vulnificus strain CG100 | GCF_002903465.1 |
| Vibrio vulnificus strain FDAARGOS_116 | GCF_001471305.2 |
| Vibrio vulnificus strain FORC_054 | GCF_002863725.1 |
| Vibrio vulnificus strain LSU2098 | GCF_002903725.1 |
| Vibrio vulnificus strain NCTC11066 | GCF_900460445.1 |
| Vibrio vulnificus strain YJ016 | GCF_000009745.1 |

**Table S2.** ***Vibrio*** **genomes used for multi-species tree (Fig. 3)**

| **Sequence Name** | **RefSeq or GenBank Accession** |
|---|---|
| Vibrio anguillarum strain JLL237 | GCF_002211985.1 |
| Vibrio anguillarum strain S2 2/9 | GCA_001989895.1 |
| Vibrio anguillarum strain VIB43 | GCF_002287545.1 |
| Vibrio cholerae strain 12129(1) | GCF_000174115.1 |
| Vibrio cholerae strain 2012Env-9 | GCF_000788715.2 |
| Vibrio cholerae strain N16961 | GCF_000006745.1 |
| Vibrio cholerae strain O395 | GCF_000021625.1 |
| Vibrio fluvialis strain 12605 | GCF_001952955.1 |
| Vibrio fluvialis strain FDAARGOS_100 | GCF_002953375.1 |
| Vibrio fluvialis strain FDAARGOS_104 | GCF_001558415.2 |
| Vibrio furnissii strain 2419-04 | GCF_009665335.1 |
| Vibrio furnissii strain FDAARGOS_777 | GCF_006364355.1 |
| Vibrio kanaloae strain 5S-149 | GCF_000272165.2 |
| Vibrio kanaloae strain 10N.222.51.B7 | GCF_005146725.1 |
| Vibrio metoecus strain 08-2459 | GCF_009665275.1 |
| Vibrio metoecus strain 2011V-1169 | GCF_009665255.1 |
| Vibrio mimicus strain 2011V-1073 | GCF_009665195.1 |
| Vibrio mimicus strain FDAARGOS_112 | GCF_001558475.2 |
| Vibrio mimicus strain SCCF01 | GCF_001767355.1 |
| Vibrio ordalii strain Q67 | GCA_002257545.1 |
| Vibrio paracholerae strain 2014V-1107 | GCF_003311945.1 |
| Vibrio paracholerae strain 2017V-1176 | GCF_003312095.1 |
| Vibrio parahaemolyticus strain 19-021-D1 | GCF_009734325.1 |
| Vibrio parahaemolyticus strain 2012AW-0154 | GCF_009665495.1 |
| Vibrio parahaemolyticus strain FDAARGOS_115 | GCF_001558495.2 |
| Vibrio scophthalmi strain VS-05 | GCF_001687805.1 |
| Vibrio scophthalmi strain VS-12 | GCF_001685465.1 |
| Vibrio vulnificus strain 93U204 | GCF_000746665.1 |
| Vibrio vulnificus strain CECT 4999 | GCF_002215135.1 |
| Vibrio vulnificus strain FORC_054 | GCF_002863725.1 |

**Table S3. Bacterial strains and plasmids.**

| Strain or Plasmid | Genotype/Description | Internal Strain Ref. | Reference and/or Source |
|---|---|---|---|
| *V. cholerae* V52 | O37 serogroup strain isolated in Sudan in 1968 from a clinical sample; $Sm^R$ | DU167 | John Mekalanos (Harvard Medical School, Boston, MA, USA) |
| *V. cholerae* V52 Δ*vasK* | V52 deleted for the T6SS membrane complex component *vasK*; $Sm^R$ | DU168 | MacIntyre *et al*., 2010 |
| *V. cholerae* DL4211 | O123 serogroup strain, isolated from the Rio Grande river delta (USA) in 2008; Aux3-naïve; $Sm^R$ | DU608 | Daniele Provanzano (UTRGV, Brownsville, TX, USA |
| *V. cholerae* DL4211 Δ*vasK* | DL4211 deleted for the T6SS membrane complex component *vasK*; $Sm^R$ | DU357 | Unterweger *et al*., 2012 |
| *V. anguillarum* VIB43 | *Vibrio anguillarum* O1 serogroup strain isolated from a diseased *Dicentrarchus labrax* (European bass) | FJS300 | Hans Rediers (KU-Leuven, Leuven, Belgium) |
| *E. coli* DH5α λpir | $F^-$ endA1 glnV44 thi-1 recA1 relA1 gyrA96 deoR nupG φ80lacZΔM15 Δ(lacZYA-argF) U169 hsdR17 ($r_K^-$ $m_K^+$) phoA, $\lambda^-$ | FJS010 | Platt *et al*., 2000 |
| *E. coli* BL21(DE3) pLysS | Str. B $F^-$ *ompT gal dcm lon* $hsdS_B(r_B^-\ m_B^-)$ λ(DE3 [*lacI lacUV5-T7p07 ind1 sam7 nin5*]) [$malB^+$]$_{K-12}$($\lambda^S$) pLysS[*T7p20* $ori_{p15A}$];$Cm^R$ | HM016 | Studier *et al.* 1986 |
| *E. coli* BL21(DE3) pLysS ; pET26b(+) ; pET22b(+) | BL21(DE3) carrying empty pET26b(+) and empty pET22b(+) expression vectors; $Cm^R$, $Kan^R$, $Amp^R$ | HM020 | This study |
| *E. coli* BL21(DE3) pLysS ; pET26b(+)-*tseL*-6xHis ; pET22b(+) | BL21(DE3) strain for the expression of TseL-6xHis ; $Cm^R$, $Kan^R$, $Amp^R$ | FJS444 | This study |
| *E. coli* BL21(DE3) pLysS ; pET26b(+) ; pET22b(+)-*tsiV1*-6xHis | BL21(DE3) strain for the expression of TsiV1-6xHis ; $Cm^R$, $Kan^R$, $Amp^R$ | FJS445 | This study |
| *E. coli* BL21(DE3) pLysS ; pET26b(+)-$Aeff^{V09}$-6xHis ; pET22b(+) | BL21(DE3) strain for the expression of $Aeff^{V09}$-6xHis ; $Cm^R$, $Kan^R$, $Amp^R$ | FJS446 | This study |
| *E. coli* BL21(DE3) pLysS ; pET26b(+) ; pET22b(+)-$Aimm^{V09}$-6xHis | BL21(DE3) strain for the expression of $Aimm^{V09}$-6xHis ; $Cm^R$, $Kan^R$, $Amp^R$ | FJS447 | This study |

| | | | |
|---|---|---|---|
| *E. coli* BL21(DE3) pLysS ; pET26b(+)-*tseL*-6xHis ; pET22b(+)-*tsiV1*-6xHis | BL21(DE3) strain for the co-expression of TseL-6xHis and TsiV1-6xHis ; $Cm^R$, $Kan^R$, $Amp^R$ | FJS448 | This study |
| *E. coli* BL21(DE3) pLysS ; pET26b(+)-*tseL*-6xHis ; pET22b(+)-$Aimm^{V09}$-6xHis | BL21(DE3) strain for the co-expression of TseL-6xHis and $Aimm^{V09}$-6xHis ; $Cm^R$, $Kan^R$, $Amp^R$ | FJS449 | This study |
| *E. coli* BL21(DE3) pLysS ; pET26b(+)-$Aeff^{V09}$-6xHis ; pET22b(+)-$Aimm^{V09}$-6xHis | BL21(DE3) strain for the co-expression of $Aeff^{V09}$-6xHis and $Aimm^{V09}$-6xHis ; $Cm^R$, $Kan^R$, $Amp^R$ | FJS450 | This study |
| *E. coli* BL21(DE3) pLysS ; pET26b(+)-$Aeff^{V09}$-6xHis ; pET22b(+)- *tsiV1*-6xHis | BL21(DE3) strain for the co-expression of $Aeff^{V09}$-6xHis and TsiV1-6xHis ; $Cm^R$, $Kan^R$, $Amp^R$ | FJS451 | This study |
| Plasmids | | | |
| pET26b(+)-*tseL*-6xHis | pET26b(+) with inserted copy of *tseL* | FJS439 | This study |
| pET22b(+)-*tsiV1*-6xHis | pET22b(+) with inserted copy of *tsiV1* | FJS442 | This study |
| pET26b(+)-$Aeff^{V09}$-6xHis | pET26b(+) with inserted copy of $Aeff^{V09}$ | FJS438 | This study |
| pET22b(+)-$Aimm^{V09}$-6xHis | pET22b(+) with inserted copy of $Aimm^{V09}$ | FJS441 | This study |

**Table S4. Primers.**

| Ref | Name | Sequence* |
|---|---|---|
| FJS328 | pET26_N16961_*tseL*_His_F | ggccatggatatcggaattaattcgATGGATTCATTTAATTATTGCGTG |
| FJS329 | pET26_N16961_*tseL*_His_R | gatctcagtggtggtggtggtggtg***tcctcctcc***TCTTATTTGCACCTTGATTTCATC |
| FJS326 | pET26_V09_Aeff_His_F | ggccatggatatcggaattaattcgATGGATTCATTTAACCATTGCG |
| FJS327 | pET26_V09_Aeff_His_R | gatctcagtggtggtggtggtggtg***tcctcctcc***TTGTAGTTGTTCCTTAATTTCATCAG |
| FJS330 | pET22_N16961_*tsiV1*_His_F | ggccatggatatcggaattaattcgATGAAGTTATTGAATAATCTTGCAATAAAAAAG |
| FJS331 | pET22_N16961_*tsiV1*_His_R | gatctcagtggtggtggtggtggtg***tcctcctcc***ATTATCATCAGATACCACTGCTG |
| FJS332 | pET22_V09_Aimm_His_F | ggccatggatatcggaattaattcgATGAAGTTATTGAATAACCTCGC |
| FJS333 | pET22_V09_Aimm_His_R | gatctcagtggtggtggtggtggtg***tcctcctcc***TTGCGCTACAGCGACTTG |
| FJS334 | pET22_pET26_insert_verification_F | TGTGAGCGGATAACAATTCCC |
| FJS335 | pET22_pET26_insert_verification_R | AGCCAACTCAGCTTCCTTTC |

* For all Gibson assembly primers, overlap regions are shown in lowercase and added glycine linker is shown in ***bold/italic.***
